# Two different and robustly modeled DNA binding modes of Competence Protein ComP - systematic modeling with AlphaFold 3, RoseTTAFold2NA, Chai-1 and re-docking in HADDOCK

**DOI:** 10.1101/2024.11.23.625027

**Authors:** Stian Aleksander Helsem, Kristian Alfsnes, Stephan A. Frye, Alexander Hesselberg Løvestad, Ole Herman Ambur

## Abstract

The competence protein ComP is a Type IV minor pilin and the extracellular DNA binding protein involved in natural transformation in the human pathogens *Neisseria gonorrhoeae*, *Neisseria meningitidis, Eikenella corrodens* and related Neisseriaceae bacteria. Details of the DNA binding mode of ComP is enigmatic, and the 3D structure of the DNA :: protein complex remains unresolved. Here we characterize the ComP orthologs in a set of Neisseriaceae family members, model their common structural domains and their interaction with different preferred 12 base pair long DNA binding motifs, DNA Uptake Sequences (DUS) and scrambled versions of these. Through systematic in silico modeling using AlphaFold 3, RoseTTAFold2NA, and Chai-1 and model comparisons, we bring a new understanding of the interactions between DNA and ComP. We report six distinct binding modes of which two, here named Epsilon and Gamma, were robustly modeled across platforms and different ComPs. The characteristics and robustness of the predicted models and DNA binding modes from each tool are assessed and discussed. This work expands the knowledge on the ComP :: DUS interaction and guides further wet- and dry-lab systematic and experimental characterization of these complexes through which molecular and clinical interventions may be developed.

## Introduction

The development of deep learning protein structure prediction algorithms is now allowing the study of protein function and interactions in a detail never seen before. AlphaFold3 (AF3) (Abramson et al., 2024), RoseTTAFold2NA (RF2NA) (Baek et al., 2024), RosETTAFold-All-Atom (Krishna et al., n.d.), NeuralPLexer (Qiao et al., 2024), DiffDock (Corso et al., 2022) and more have led the way to model protein interactions with other proteins, ligands and nucleic acids. Recently Chai-1 was released based with single model training and new functionality (Chai Discovery, 2024) and new implementations of AlphaFold 3 (AF3) are continuously being developed, with the release of HelixFold3 (Team, 2024) and Boltz-1 (Wohlwend et al., 2024) and the field is rapidly advancing. The underlying platform algorithms differ in fundamental ways and therefore also their emphasis on various confidence metrices. The Predicted Aligned Error (PAE) was introduced with AlphaFold and AF3 places a strong emphasis on PAE for providing information regarding confidence of predicted spatial relationships between residues and is used together with different Local Predicted Confidence Difference Test (LDDT) scores (Abramson et al., 2024). PAE is reported by default in AF3. RF2NA filters models based on a PAE < 10 threshold as acceptable but does not emphasize PAE-resolution below this this cut-off and like AF3, reports confidence from Local Distance Difference Test (lDDT) relative to available experimental structures and Predicted Local Distance Difference Test (pLDDT) as the model-intrinsic confidence score at the residue level (Baek et al., 2023). Chai-1 also applies per-residue and chain interface confidence metrics (LDDT and iLDDT) yet also integrates whole structure reliability from DockQ scoring (Chai discovery, 2024). In having an emphasis on PAE, AF3 also applies a Chain Pair PAE Minimum (CPPM) metric to assess the confidence of the interaction between two chains, such as DNA and protein, within the complex and propose the metric useful to distinguish binding from non-binding interactions (Alpha Fold Server). Here, we explore the hypothesis that using fundamentally different platforms to model the structure of an unresolved DNA :: Protein complex may educate our understanding of their specific interaction and potentially provide a modeling consensus.

Natural transformation is the uptake and recombination of DNA and is an evolved mechanism of many Gram positive and Gram negative bacteria (reviewed in Dubnau & Blokesch, 2019). Competent bacteria able to use natural transformation to acquire DNA display a range of different adaptations and competence states. Horizontal gene transfer mechanisms facilitate the spread of advantageous traits including antibiotic resistance and virulence genes and are therefore of clinical interest (Arnold et al., 2022).Transformation in the Neisseriaceae family containing human pathogens *Neisseria gonorrhoeae*, *Neisseria meningitidis* and *Eikenella corrodens* together with opportunistic and commensal relatives is enhanced when short 10-12 nucleotide DNA binding motifs (e.g. ATGCCGT<u>CTG</u>AA in *N. meningitidis/ gonorrhoeae*) are present in the transforming DNA (reviewed in Ambur et al., 2016). These motifs are named DNA Uptake Sequences (DUS) and are employed as standard practice for genetic manipulation of *Neisseria* species (Dillard & Chan 2024), for rapid detection of *N. gonorrhoeae* using DUS-targeting gold nanoparticles (Carter et al., 2022) and for molecular typing of *Kingella kingae* (Basmaci et al., 2014). DUS are genomically enriched in permissive parts of their respective genomes (Treangen et al., 2008) and shown *in vivo* to facilitate DNA mobility within species or between closely related genera (Frye et al., 2013). Extracellular DNA containing multiple DUS has been shown trapped at the peripheral edges of gonococcal colonies where it readily transforms (Bender et al, 2022). *In vitro* acrylamide EMSA and surface plasmon resonance experiments using purified ComP, DUS-DNA and competitive DNA have shown that ComP preferentially binds it cognate DUS with high affinity and that this is an inherent property of ComP homologs (Cehovin et al., 2013; Berry et al., 2013; 2016). Eight dialects of DUS have been described, all of which have a conserved Watson strand CTG core essential for DUS-mediated transformation by means of the competence protein ComP (Frye et al., 2013; Berry et al., 2013). In an evolutionary perspective, DUS provides means to acquire homologous DNA for allelic re-shuffles (Ambur et al., 2016).

ComP is a minor type IV pilin first identified as a positive effector of DUS-mediated transformation in *N. gonorrhoeae* (Aas et al., 2002) and later identified as the DNA-binding DUS receptor in *Neisseria subflava* and other *Neisseria sp.* (Cehovin et al., 2013). *In vivo* overexpression of *N. subflava* ComP in a *N. meningitidis* model has been shown to increase transformation also in a DUS-independent manner almost up to the level of DUS-specific transformation (Berry et al., 2013). At least two different DNA binding modes of ComP may therefore exist, one DUS-specific and one or more unspecific. Alternatively, DUS-specific and unspecific DNA binding may share the same singular binding mode assuming only weak DUS sequence binding biases with more relaxed interactions to parts of the DUS outside the essential CTG core in accordance with wet-lab experiments (Berry et al., 2013; Frye et al., 2013). ComP is integrated into the type IV pilus (T4P) protruding from the neisserial outer membrane together with the major pilin PilE and other minor pilins (Berry et al., 2016). All the minor pilins of T4P are structurally similar at the α-helical N-terminus required for pilin multimer assembly (reviewed in Jacobsen et al., 2020). Relative to other minor pilins, ComP has a unique, complex and exposed C-terminal globular domain where DNA can bind. The high-resolution 3D structures of ComP (*N. meningitidis* and *N. subflava*) have been reported and are based on crystallography of soluble ComP-maltose-binding-protein fusion proteins and NMR of 6-His tagged ComP (Berry et al., 2016). ComP_N.men_ and ComP_N.sub_ are structurally very similar and are characterized by an N-terminal α-helix, five antiparallel β-strands, the α1-β1- and β1-β2-loops, a DD-region, and two disulfide bridges (Cys-Cys). Intramolecular disulfide bridges are formed in exported proteins by thiol-disulfide oxidoreductases (DsbAs) and wet-lab experiments have shown that *N. meningitidis* DsbA knock-out mutants have reduced DNA uptake in a DUS-dependent manner (Sinha et. al, 2008). Whether this phenotype could directly link to ComP is unknown. The two ComP disulfide bridges staples the DD-loop to the β1-strand and the β4-β5 strands together, were found crucial for DUS-binding in electrophoretic mobility shift assays (Cehovin et al., 2013). ComP has an electropositive stripe, and individual substitutions of three positively charged amino acids (R78A, K94A, and K108A) were shown *in vivo* to affect binding and/or transformation negatively (Cehovin et al., 2013). Several Neisseriaceae have been demonstrated to have higher transformability and binding affinity for DUS containing DNA compared to other sequences using transformation experiments and biophysical approaches (Goodman & Scocca, 1988; Aas et al., 2002, Frye et al., 2013; Berry et al., 2013, 2016; Cehovin et al., 2013). NMR spectra showed that the DD-region, the tip of the ß1-ß2 loop and part of the α1-ß1 loop were involved in DUS binding and were modeled to establish contacts with bases in successive grooves of the dsDNA (Berry et al., 2016). ComP has been found to preferentially mediate transformation of dsDNA, although single-stranded Crick strand DUS (TTCAGACGGCAT) shows some transforming ability in a *N. gonorrhoeae* model relative to no-DUS and the Watson strand DUS (ATGCCGTCTGAA) (Duffin & Seifert, 2012). Another study exploring uptake and transformation of fluorescently labelled ssDNA and dsDNA found similarly that efficient DUS recognition was dependent on double-stranded DNA whereas the rates of transport across the plasma membrane were comparable (Hepp et al., 2016).

The ComP 3D structure with bound DUS/DNA remains unresolved. All major structural work has focused on the ComP proteins of *N. meningitidis* and *N. subflava* (Berry et al., 2013, 2016; Cehovin et al., 2013). Fewer efforts have been put into studying the ComP :: DUS/DNA binding mode in silico. Berry et al. (2016) and Hughes-Games (2020) performed docking analyses on *N. subflava* NJ9703_ComP_ (here N.sub_nat_) and *N. meningitidis* 8013_ComP_ (here N.men_nat_). Their molecular docking results both seem to show (our interpretation) that the α1-β1 and β1-β2 loops interact with major groove (MG) and the DD-region with minor groove (mG) of the DNA.

No systematic modeling of ComP with DUS/DNA has been performed using the now available and fundamentally different deep learning-based platforms for structure prediction of such complexes. AF3, RF2NA and Chai-1 represent the state-of-the-art of modeling protein-nucleic acid (protein-NA) complexes and may hence provide means to better understand the ComP :: DUS/DNA interaction. Using five of the DUS dialects from Frye et al. (2013) as the basis for comparatively modeling ComP :: DUS/DNA interactions using different modeling algorithms, this study brings a new understanding of the ComP :: DUS/DNA complex and characterize two different and robust DNA binding consensuses.

## Materials and methods

### Data material

ComP orthologs were obtained using a series of BLASTp searches (Camacho et al., 2009), using the structures with PDB accession 2NBA (Berry et al., 2013), 2M3K (Cehovin et al., 2013) and 5HZ7 (Berry et al., 2013) as queries against the RefSeq non-redundant database in the NCBI web server, specifying the Neisseriaceae clade as target. GenBank/RefSeq ComP primary structure and accession numbers are listed in Table S1. Trimming of the primary ComP structure removed 28 amino acids of the N-terminal part of the mature ComP_Nmen_ and the equivalent sections in the other six orthologous ComPs as they were trimmed in the published 3D structures (Cehovin et al., 2013; Berry et al., 2013). ComP primary structures from seven Neisseriaceae species were modeled with dsDNA containing their native 12 bp DUS dialects in this study (Table S1). Native DUS dialects refer to the genomically most abundant 12-mer DUS of each species as identified and described previously (Ambur et al., 2007; Frye et al., 2013). The ComP :: DUS pairs were: *Eikenella corrodens* ATCC 23834 ComP :: AG-eikDUS (E. cor_nat_), *Neisseria meningitidis* MC58 ComP :: AT-DUS (N. men_nat_), *Kingella denitrificans* ATCC 33394 ComP :: AA-king3DUS (K. den_nat_), *Neisseria mucosa* ATCC 25996 ComP :: AG-mucDUS (N. muc_nat_), *Bergeriella denitrificans* ATCC 33394 ComP :: AG-DUS (B. den_nat_), *Neisseria cinerea* ATCC 14685 ComP :: AT-DUS (N. cin_nat_) and *Neisseria subflava* NJ9703 ComP :: AG-DUS (N. sub_nat_). These native ComP :: DUS modeling complexes are described with the suffix “nat” appended to the species name (e.g. N. sub_nat_). Equivalently when using scrambled native DUS in ComP :: DNA modeling the suffix “scr” was appended to the species name (e.g. N.sub_scr_).

### Modeling of ComP_nat_s

To model ComP_nat_s for the first time, seven different ComPs were modeled with their native 12 bp DUS (dsDNA) in three different deep-learning structure prediction platforms: AF3 (web server), RF2NA (local installation) and Chai-1 v. 0.2.0 (local installation). The flowchart followed is presented in Fig 1.

**Figure 1.**
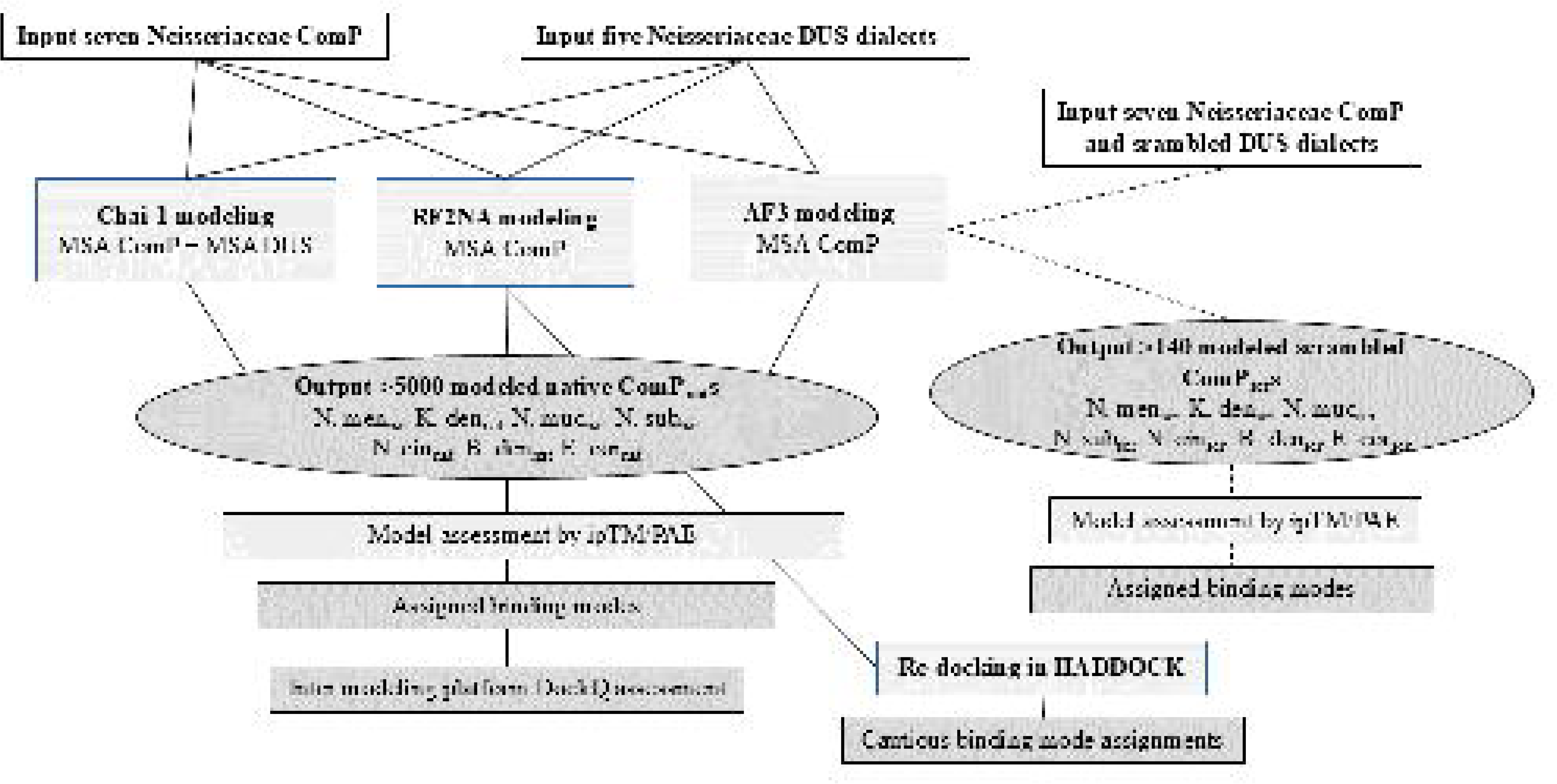
Flowchart of the study showing the steps taken to model native ComP_nat_ with Chai-1, RF2NA and AF3 and re-docking of top-ranking RF2NA models in HADDOCK. AF3 modeling of ComP_scr_ follow dashed lines on the right side. Boxes with inputs are white, processes light gray and outputs dark grey. All data sources and parameters used are described in the text.

A minimum of 20 native AF3 replicates (33 for B. den_nat_ and 37 for N. sub_nat_) and 100 native runs with different seeds in RF2NA and Chai-1 were performed for each ComP_nat_. Customization of multiple sequence alignment (MSA) input to Chai-1 allowed for applying also an MSA of DUS in addition to the ComP MSA, the details of the different MSA schemes explored are listed in Table S2. These MSAs were generated using HHblits v. 3.0.0 (Steinegger, Meier, et al., 2019) to search the Big Fantastic Database (BFD) (Jumper et al., 2021; Steinegger, Mirdita, et al., 2019; Steinegger & Söding, 2018) and Uniref30 (Mirdita et al., 2022) databases for ComP MSAs, using the same HHblits parameters as in RF2NA. DUS MSAs were created based on genomic DUS counts in the seven selected species (as in Frye et al., 2013). 10 replicates were run for each MSA scheme. This generated a grand total of 3500 mmCIF/PDB models in Chai-1 (500 per ComP_nat_), 850 models in AF3, with more than 100 models per ComP_nat,_ and 700 models in RF2NA, with 100 models per ComP_nat_. Criteria for assigning successful models were ipTM > 0.6 in AF3 and Chai-1 and PAE < 10 in RF2NA, following cut-off recommendations on the AF3 web server for ipTM and in Baek et al. (2024) for PAE (Table 1).

**Table 1.** Model quality measures and confidence metrices used to assess predicted models from Chai-1, AF3, RF2NA and HADDOCK.

| Metrics | Score/test description | Range | Unit | Applied and reported | Relevant platform(s) |
| --- | --- | --- | --- | --- | --- |
| ipTM | Predicted interface template modeling score | 0-1<br>Poor-excellent | - | $\geq 0.6$<br>considered successful | AF3<br>Chai-1 |
| PAE | Predicted aligned error | Continuous | Å | $< 10$<br>considered successful in RF2NA | AF3<br>Chai-1<br>RF2NA |
| pLDDT | Predicted local distance difference test | 0-100<br>Very poor-very high confidence | - | Significant differences between models | AF3<br>Chai-1<br>RF2NA |
| CPPM* | Chain pair PAE minimum | Continuous | Å | Significant differences between models | AF3 |
| HADDOCK score | Weighted sum of various energy terms | Continuous | - | Normal mode | HADDOCK |
| DockQ** | Quality measure of similarity | Continuous | - | 0-1<br>Poor-excellent | AF3, Chai-1 and RF2NA |

\* Used here to distinguish probable binders from non-binders in interactions between chains in AF3
\*\* Used here to measure consistency and mode convergence in predicted $\text{Comp}_{\text{nat}}$ models within and between AF3, Chai-1 and RF2NA (Basu & Wallner, 2016; Mirabello & Wallner, 2024).

To investigate the confidence of AF3 to model the seven ComP_nat_s in a DUS-specific manner as measured by CPPM, AF3 modeling using also scrambled DUS sequences was carried out. All seven ComPs were modeled with scrambled 12 bp versions of their native DUS dialects in AF3, not allowing the scrambled DNA strings to contain the conserved trinucleotide 5’-CTG-3’ nor tetra-homomers (i.e. “GGGG”) in the sequence, while containing the same nucleotide content as the native DUS. Wilcoxon rank sum significance tests were then performed to determine whether runs with native DUS yielded higher confidence metrics (see below) than scrambled DUS runs. All 140 AF3 generated ComP_scr_s models were assigned binding mode as for ComP_nat_s described below.

To evaluate the modeling reproducibility of the different binding modes while also assessing the docking approaches used in previous studies (Berry et al., 2016; Hughes-Games, 2020), re-docking experiments of RF2NA outputs were conducted in HADDOCK (Teixeira et al., 2024). The top-ranking RF2NA ComP_nat_s were re-docked. Following the suggestion by (Esmaeeli et al., 2023), prior to molecular docking in HADDOCK, the first 10 all-atom non-trivial normal modes with lowest frequency for both the ComPs and DUS were generated using the R module bio3d (Grant et al., 2006, 2021; Skjærven et al., 2014; 2016). Selected normal mode conformations were fed into HADDOCK as ensemble protein and DNA PDB files and cross-docked (all against all), generating 400.000 rigid bodies per ComP_nat_, of which, after flexible and molecular dynamics refinement, the top 200 models (HADDOCK score) were assigned binding mode. Because the input DNAs were highly distorted (due to normal mode calculation), it was often challenging to discern DNA grooves (mG and MG) and assign binding mode in the re-docking outputs. Binding modes for all HADDOCK models were therefore very cautiously considered. This was also why the HADDOCK output models were not used in the cross-platform DockQ consistency analyses. For N. sub_nat_, all solvent-accessible ComP residues that showed an NMR CSP shift below DNA concentration of 20 mM in Berry et al., 2016 were specified as ambiguous restraints. For the other six ComP_nat_s, the same residues were used as ambiguous restraints only if conserved in the ComP alignment (Fig S2).

All Chai-1 and AF3 models having ipTM ≥ 0.6 and RF2NA models having PAE < 10 (ipTM is not an output in RF2NA) were deemed robust and inspected in 3D using PyMOL v. 3.0.0 and binding mode assigned according to the descriptions below. In ComP_nat/scr_ models for which ipTM never reached the 0.6 cut-off, all models were inspected in 3D and assigned binding mode. Based on similarity across models in how central protein domains and DNA grooves interact, general binding modes were defined (Table 2).

**Table 2.** ComP DNA binding modes. Modes are differentiated and assigned by how four central ComP protein domains interact and enter the DNA minor-(mG) and major (MG) grooves of DNA.

| Comp domain/<br>Binding mode | $\alpha 1$-$\beta 1$ loop | $\beta 1$-$\beta 2$ loop | DD-region | $\beta 4$- and $\beta 5$-<br>sheets | Comment |
| --- | --- | --- | --- | --- | --- |
| $\gamma$ (Gamma) | mG | mG | MG | | |
| $\delta$ (Delta) | mG | mG | mG | | |
| $\epsilon$ (Epsilon) | MG | MG | mG | | |
| $\zeta$ (Zeta) | MG | MG | MG | | |
| $\eta$ (Eta)* | | | | MG | N-terminus<br>and DNA<br>end close |
| $\theta$ (Theta)** | MG | MG | MG | | DNA partly<br>unwound |
| o (Other) | - | - | - | - | Unlike rest |

\* Only in HADDOCK for K. den<sub>nat</sub>, B. den<sub>nat</sub> and N. men<sub>nat</sub>. $\eta$ part of o in Table 3.
\*\*Only in Chai-1 for N. cin<sub>nat</sub>

Representative examples of ComP_nat_ complexes in the six different binding modes Gamma, Delta, Epsilon, Zeta, Eta and Theta are shown in Fig S1 and representative PAE-plots for each ComP_nat_ assigned modes in Fig S4-S6. Since PAE plots were not generated by default in either Chai-1 nor RF2NA, these were made using customized scripts.

### Internal consistency and cross-platform convergence

To investigate intra-platform consistency of AF3, Chai-1 and RF2NA results separately, all ComP_nat_ against all ComP_nat_ DockQ scores were computed for the top-ranking models across replicate runs within the same platform. This was devised as a test of platform consistency for repeatedly predicting the same binding mode. Additionally, the degree of agreement in the results across AF3, Chai-1 and RF2NA was quantified by comparing the DockQ scores for all robust models against all other robust models. To avoid giving weights to lower ranking models, only AF3 and Chai-1 models having ipTM ≥ 0.6 were considered robust in the cross-platform DockQ comparisons as advised (Abramson et al., 2024; Zhang & Skolnick, 2004), except for N. men_nat_ and N. cin_nat_ in AF3 and N. muc_nat_ in Chai-1, for which ipTM ≥ 0.6 was never reached and all models were included in these cases. Likewise, since ipTM is not reported in RF2NA only those RF2NA models having mean interface PAE < 10 were considered robust and included in the assessment as advised by Baek et al., 2024.

## Results

### The conserved ComP structure

Here, using AF3, the published structures of two ComP proteins (ComP_N.men_ and ComP_N.sub_) (Berry et al., 2016; Cehovin et al., 2013) and five additional orthologous proteins from other Neisseriaceae members (*B. denitrificans*, *E. corrodens*, *N. mucosa*, *N. cinerea* and *K. denitrificans*) were aligned and compared (Fig 2).

**Figure 2.**
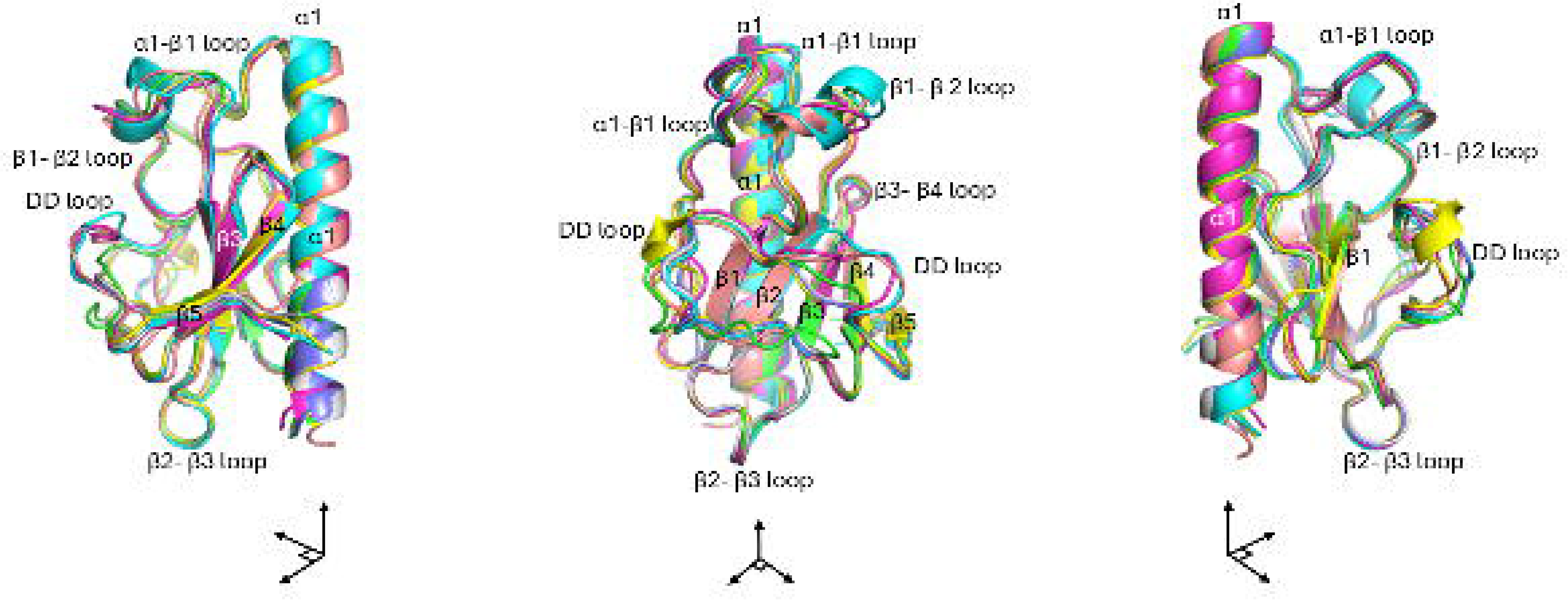
Structural overlay of ComPs from seven Neisseriaceae species. These are AF3 models aligned on the α-carbons of the model with the best ipTM score, Purple: ComP_K.den_; Green: ComP_N.sub_. Turquoise: ComP_B.den_. Yellow: ComP_N.muc_. Pink: ComP_E.cor_. Grey: ComP_N.men_. Blue: ComP_N.cin_. The β-strands are numbered, and domains labelled. The three representations are 90° right turns around the N-terminal α-helix aligned on the y-axis.

The structural alignment of the AF3 models revealed a high degree of similarity across all seven ComPs and that all contained the same domains as previously described (Berry et al., 2016; Cehovin et al., 2013). Identifiable and ubiquitous domains are the N-terminal α-helix, five antiparallel β-strands, β-strand connecting loops, the DD-region, and two essential disulfide bridges (connecting β1-DD-loop and β4-β5) (Berry et al., 2016). Minor differences were seen across the ComPs, including prediction of a short, additional α-turn embedded in the β1-β2-loop in ComPs of *B. denitrificans* and *E. corrodens* and a short α-turn located in the DD-region in ComP of *N. mucosa*.

Minor structural variations in different domains for the same ComP were observed across different replicate runs, even within the same platform. Particularly, parts of the DD-region and the β1-β2 loop of ComP were modelled generally less consistently. In the modeled structures in Fig 2, these flexible regions can be seen as imperfectly aligned domains (separated protein strands). These are located in indel-variable regions of the investigated ComPs, as shown in their aligned primary structure (Fig S2).

Due to strong overall structural predicted consistency across platforms, ComP structure overlays from Chai-1 and RF2NA are not shown beyond their respective ComP_nat_s in Figure 3 below, and their representative ComP_nat_ PAE plots in the supporting data (Figs S4-S7). We note however that despite overall structural consistency, RF2NA generally produced models lacking the disulfide bridge connecting the β1-strand and the DD-region even though these were present in all three experimentally predicted template structures commonly used in the MSAs (2NBA, 5HZ7 and 2M3K). Furthermore, a smaller fraction of the RF2NA models, also the disulfide bridge connecting β4- and β5-sheets were lacking which could affect the integrity of the β-sheet. In contrast to all AF3 and Chai-1 models, RF2NA was therefore found less prone to maintain modeled and functionally essential disulfide bridges in ComP.

**Figure 3.**
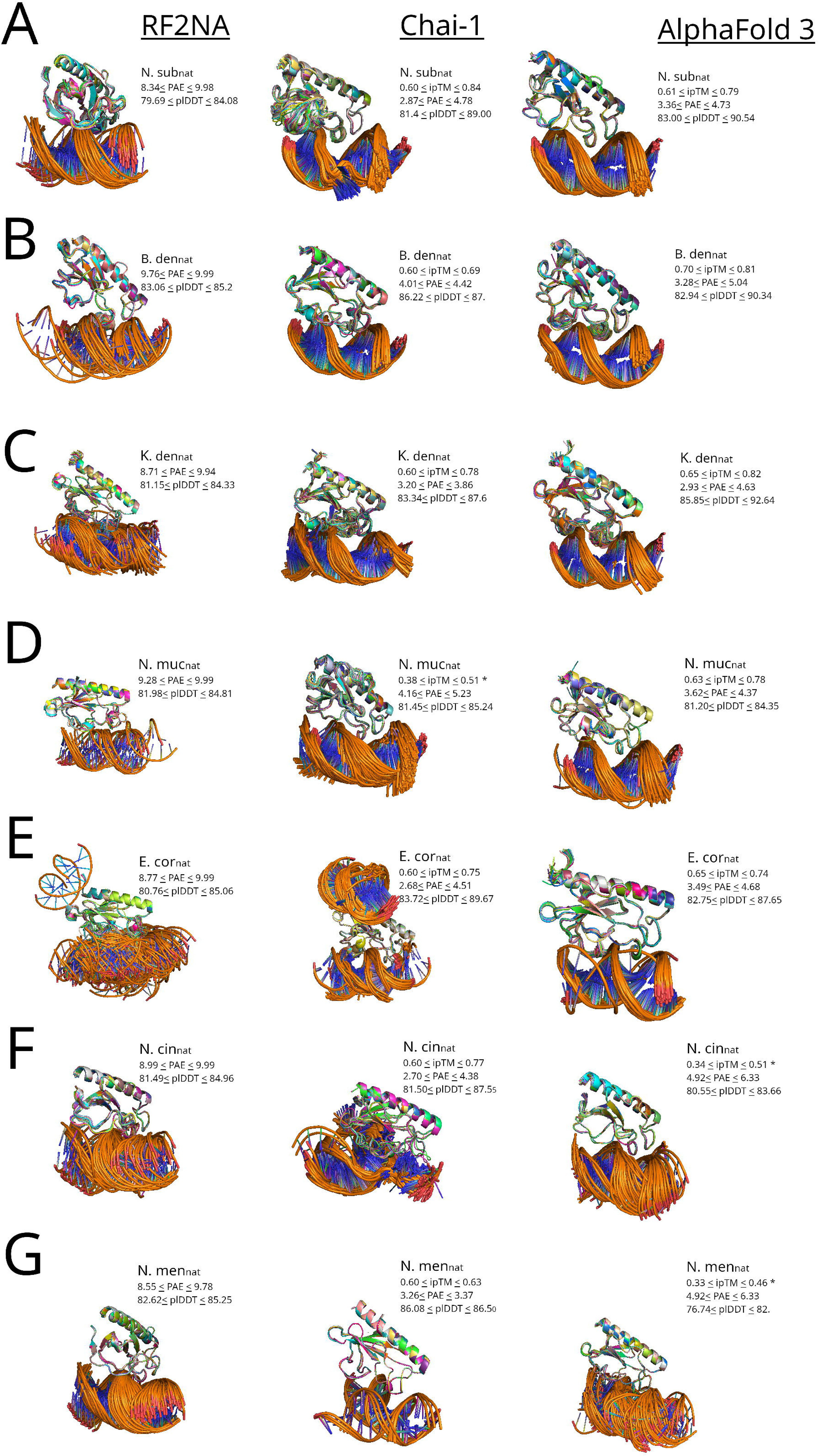
Robust ComP_nat_ models from RF2NA, Chai-1 and AF3 for all seven ComP_nat_ complexes (A-G). **A**: N.sub_nat_; **B**: B.den_nat_; **C**: K. den_nat_; **D**: N. muc_nat_; **E**: E.cor_nat_; **F**: N. cin_nat_; and **G**: N. men_nat_. PAE and pLDDT score ranges of models included in the overlay are listed next to each ComP_nat_ overlay and in addition ipTM score ranges for AF3 and Chai-1 models. Each overlay is aligned on the α-carbon of the models with the best ipTM score or lowest PAE for RF2NA models. For each ComP_nat_ modeled in AF3 and Chai-1, only the subset of models with ipTM ≥ 0.6 are represented and when no models were predicted above the ipTM threshold (Fig 3 F, G and D and Fig 6 A), all predicted models are represented and labeled with *. For RF2NA, only models having PAE ≤10 are represented.

### Modeling and re-docking results

A series of iterated modelling experiments of all seven ComP_nat_s were conducted in all three platforms, RF2NA, Chai-1 and AF3 to provide a grand total of 3620 ComP_nat_ model complexes. These ComP_nat_ models are shown in Fig 3 as platform specific complex overlays and their binding modes are detailed in Table 3 together with 1386 models from the re-docking in HADDOCK.

**Table 3.** Total counts of ComP_nat_ models from all three platforms AF3, Chai-1, RF2NA and re-docking in HADDOCK with assigned binding modes. Greek letter symbols for modes are ⍰ = Epsilon, ***ζ*** = Zeta, ***γ*** =Gamma, ***δ***=Delta and **o**=other (here incl. Eta and Theta)

| Tool | AF3 |  |  |  |  | Chai-1 |  |  |  |  | RF2NA* |  |  |  |  | HADDOCK |  |  |  |  |
| --- | --- | --- | --- | --- | --- | --- | --- | --- | --- | --- | --- | --- | --- | --- | --- | --- | --- | --- | --- | --- |
| Mode/<br>Comp<sub>nat</sub> | ⌈ | ζ | γ | δ | o | ⌈ | ζ | γ | δ | o | ⌈ | ζ | γ | δ | o | ⌈ | ζ | γ | δ | o |
| N.sub<sub>nat</sub> | 258 | - | - | - | - | 428 | - | - | - | - | - | - | 1 | 45 | - | 26 | - | 159 | - | 15 |
| B.den<sub>nat</sub> | 165 | - | - | - | - | 91 | - | - | - | - | 20 | - | 1 | - | 1 | 145 | - | 1 | - | 54 |
| K.den<sub>nat</sub> | 146 | - | - | - | - | 445 | - | - | - | - | 25 | 69 | 4 | - | - | 200 | - | - | - | - |
| N.muc<sub>nat</sub> | 105 | - | - | - | - | *50<br>0 | - | - | - | - | - | - | 10 | - | 1 | 145 | - | 53 | - | 2 |
| E.cor<sub>nat</sub> | - | - | 95 | - | - | 157 | - | 138 | - | 86 | 23 | 25 | 26 | 1 | 6 | ‡17<br>0 | ‡18 | - | - | - |
| N.cin<sub>nat</sub> | 4 | - | *96 | - | - | - | - | - | - | †395 | - | - | 40 | 12 | - | 79 | - | 121 | - | - |
| N.men<sub>nat</sub> | *21 | - | *76 | *3 | - | 12 | 1 | - | - | - | - | - | 56 | 35 | - | 130 | 2 | 68 | - | - |
| SUM | 699 | 0 | 267 | 3 | 0 | 163<br>3 | 1 | 138 | - | 481 | 68 | 94 | 136 | 93 | 7 | 895 | 20 | 402 | - | 69 |

\*ipTM < 0.6 or ipTM not available (RF2NA)
† All in Theta mode
‡ For E. cor<sub>nat</sub> in HADDOCK, 200 docking solutions was not reached and 188 final docking solutions were found

To investigate the modeling quality of each ComP_nat_ prediction more closely, the PAE and pLDDT distributions of each modeled complex by RF2NA, AF3 and Chai-1 are shown as scatter plots per mode and platform in Fig 4A and subdivisions of the scatters within the dominating Epsilon and Gamma modes per ComP_nat_ in the B and C panels, respectively.

**Figure 4.**
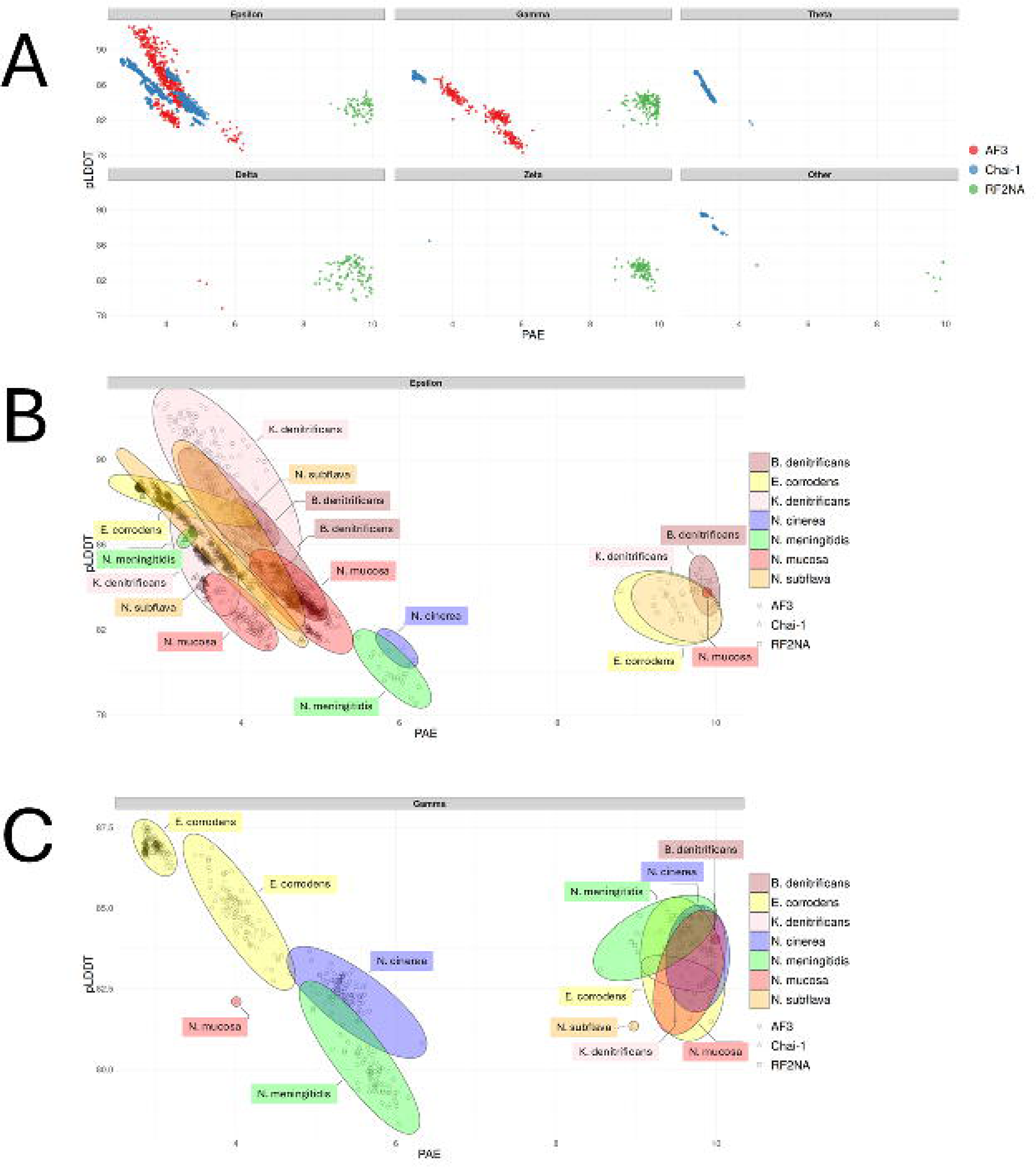
**A**. PAE/pLDDT scatter plots or all binding modes of all ComP_nat_s from AF3 (red), Chai-1 (blue) and RF2NA (green). **B**. Venn diagram of the resolved Epsilon scatters in A per color coded ComP_nat_ species. Modeling platforms are represented with symbols as assigned in the right panel. **C**. Venn diagram of the resolved Gamma models in A per color coded ComP_nat_ species. Modeling platforms are represented with symbols as assigned in right panel.

To further systematically explore AF3, Chai-1 and RF2NA mode convergence and quality of each ComP_nat_, all models with assigned modes from all platforms were paired and compared directly using DockQ scores. The DockQ scores were categorized as higher or lower according to the 0.23 (low), 0.49 (intermediate) and 0.8 (high) limits and the results are shown as violin plots in Figure 5.

**Figure 5.**
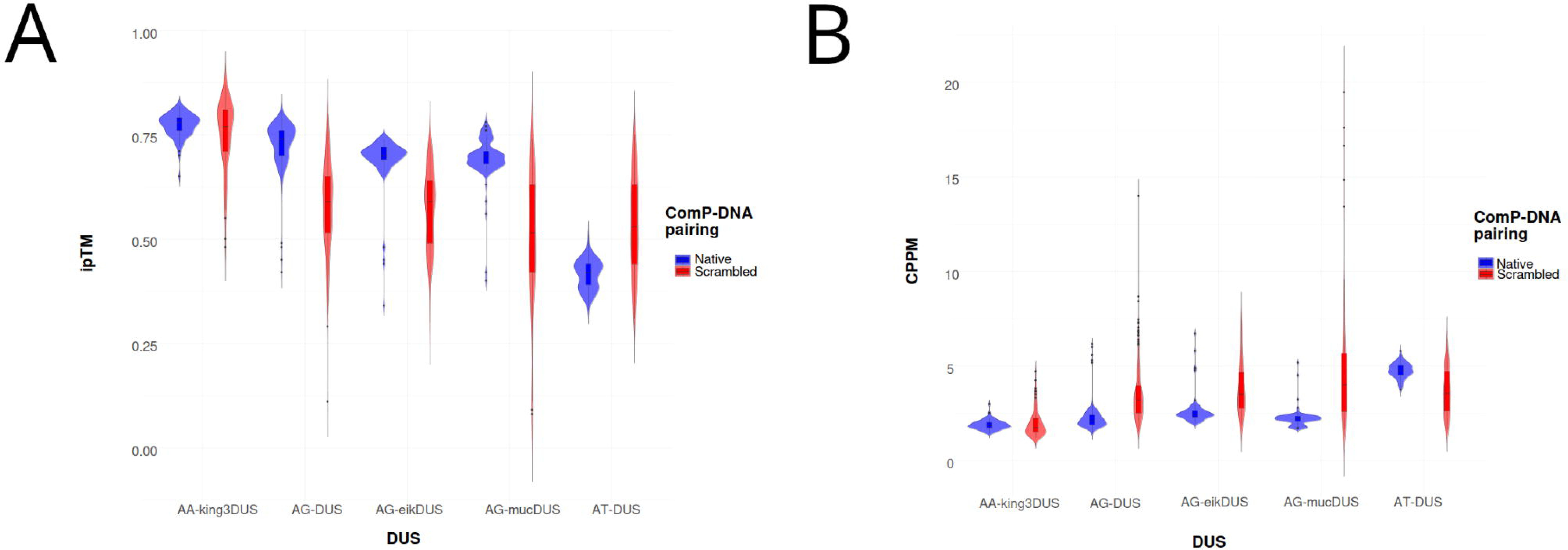
Violin plots of DockQ scores for paired all against all model comparisons of each ComP_nat_ across platforms. The median and interquartile ranges are shown as horizontal bars, and paired model comparisons are depicted as swarms inside the violins. Paired inter-platform models which were assigned to two different binding modes are shown as “Different”. Horizontal red, yellow and green dashed lines represent DockQ of 0.23 (low), 0.49 (intermediate) and 0.8 (high), respectively. Red violins represent inter-platform model pairs in Epsilon, blue Gamma, light grey Other, dark grey Delta and green Different.

### General modeling performance

The distribution of AF3 models across modes covered the widest quality PAE/pLDDT score ranges of the three platforms with PAE range 3-6 Å and pLDDT 78 (lowest)–93 (highest) (Fig 4A). The distribution of Chai-1 models covered similar yet narrower ranges as AF3 with PAE range 2.6 (lowest)–4.5 and pLDDT range 82-89. The RF2NA distributions showed a systematically higher PAE range, 8-10, than AF3 and Chai-1 across modes, whilst the pLDDT range 80-86 overlapped those of the other two platforms. The highly divergent PAE distributions show how confidence from PAE is differentially emphasized in RF2NA relative to AF3 and Chai-1. PAE, by not representing a universal metric were therefore very cautiously considered. PAE may therefore be better suited for intra-platform confidence scoring than between RF2NA and the other two platforms or in combination with other metrics such as pLLDT and DockQ.

When assessing the internal model consistency of AF3, Chai-1 and RF2NA ComP_nat_s using DockQ scores, both AF3 and Chai-1 showed higher model consistency across replicate runs of all ComP_nat_s than RF2NA. AF3 showed a significantly higher median DockQ score (0.841) compared to the other two platforms (Chai-1: 0.680; RF2NA: 0.617), as determined by pairwise Wilcoxon rank-sum tests (Code Output S3). Also, Chai-1 showed a significantly higher median DockQ score compared to RF2NA.

### The DNA binding modes

The only two modes predicted by all three platforms and in the re-docking experiments in HADDOCK were Epsilon (3295/5005) and Gamma (953/5005) (Table 3, Fig S1). Overall, AF3, Chai-1 and HADDOCK predicted more Epsilon and RF2NA more Gamma. N.sub_nat_, B.den_nat_, K.den_nat_ and N.muc_nat_ were more consistently predicted in Epsilon and E.cor_nat_, N.cin_nat_ and N.men_nat_ in Gamma. The Zeta mode (115/5005), which resembles Epsilon (Table 2), was not predicted by AF3 and mostly by RF2NA. The Delta mode (93/5005), which resembles Gamma, was not predicted by Chai-1 and mostly by RF2NA. “Other” modes were not predicted by AF3. The unique Theta mode (359/5005) was only predicted by Chai-1, which permits entry of additional MSA DNA schemes. HADDOCK also found strong general support for the Epsilon and Gamma modes and deviate from the three modeling platforms most notably in assigning Gamma for N.sub_nat_ (only 1 recorded in RF2NA), a strong Epsilon bias for E. cor_nat_ and a bias for Epsilon for N. men_nat_.

All seven ComP_nat_s were modeled in Epsilon in either AF3, Chai-1 or both. The AF3 and Chai-1 models formed clusters of models with PAE range 2.5-6 Å and pLDDT range 78-93 (Fig 4B). The AF3 Epsilon models covered a wider range (strongest and weakest) than Chai-1 Epsilon regarding both measures and included more ComP_nat_s. Fig 4B shows an overlapping region where models from four ComP_nat_s (N.sub_nat_, B.den_nat_, K.den_nat_, E.cor_nat_) converge on highly similar complexes modeled by either AF3 and Chai-1 or both (B.den_nat_). All RF2NA Epsilon models (E.cor_nat_, B.den_nat_, N.muc_nat_ and K.den_nat_) clustered together and were skewed towards considerably higher PAE range 8.5-10 than AF3 and Chai-1 and covered a pLDDT range of 82-85.

All seven ComP_nat_ were modeled in Gamma. Gamma found its strongest models for E.cor_nat_ modeled by both Chai-1 and AF3 (Fig 4B). As with Epsilon described above, the AF3 Gamma models covered a wider range than Chai-1 regarding both measures and reflected an additional three ComP_nat_ (N. cin_nat_, N. men_nat_ and a single N.muc_nat_ model). RF2NA Gamma models were clustered within narrower PAE and pLDDT ranges than AF3 and Chai-1 and were again systematically skewed toward higher PAE.

Theta models were only found for N.cin_nat_ in Chai-1 with PAE 2.5 and pLDDT 87. Theta models formed an elongated cluster along both axes (Fig 4 A) in a similar manner as the Chai-1 N.sub_nat_ Epsilon models.

The Delta mode, which resembles Gamma (Table 2), was found in only three models in AF3 (N.men_nat_), having PAE range 5-5.5 and comparatively low pLDDT range 78-82 and not in Chai-1. This mode was however, modeled repeatedly in RF2NA with PAE range 8.3-10 and pLDDT range 79-85 reflecting models for N.cin_nat_, N.men_nat_ and N.sub_nat_.

Zeta was observed once in Chai-1 (N.men_nat_) with PAE 3.5 and pLDDT 86 and repeatedly in RF2NA (K.den_nat_ and E.cor_nat_) with PAE range 8.3-10 and pLDDT range 81-84. These robust “Other” models which did not clearly fall into any of the other categorized binding modes still had high pLDDT of 88 and low PAE of 3, while those from RF2NA were again systematically skewed with PAE just below the cut-off, 9.5, and pLDDT 84.

All but one (B.den_nat_) “Other” models in Chai-1 and RF2NA were E. cor_nat_ models. The Chai-1 models were again considerably better than those of RF2NA in terms of PAE/pLDDT.

Since no ComP_nat_ modeled to the same distribution of modes across platforms, the main results, confidence scores and cross-platform DockQ consistencies as presented in Figs 2-5 and Table 3 are described in further detail below for each ComP_nat_. The statistics applied for calculating significant differences in quality scores for each ComP_nat_ are described in the Methods section and results detailed in Code output S1-3.

### Quality of the ComP_nat_s

#### N.sub_nat_

All models by AF3 (158/158) and Chai-1 (428/428) were in Epsilon, while RF2NA models were in contrast assigned Delta (45/46) and Gamma (1/46). Overall, the Chai-1 models were found to be significantly better than AF3 models and both Chai-1’s and AF3’s models were significantly better than RF2NA’s in terms of PAE and pLDDT. In the PAE/pLDDT scatter plot a smaller fraction of N.sub_nat_ models from AF3 and Chai-1overlapped each other in Epsilon and N.sub_nat_ overlapped all other ComP_nat_s except N.cin_nat_ in Epsilon and none in Gamma (Figs 4B and C). N.sub_nat_ models did however overlap in terms of PAE/pLDDT with E.cor_nat_, N.men_nat_ and N.cin_nat_ models in Delta (resolved scatter not shown). A consistently flipped-out base and DNA strand deformation were observed in the Chai-1 models (Fig 3A), showcasing the platform’s unique ability to strongly alter the DNA structure of these short templates, possibly to induce an optimized fit. Re-docking in HADDOCK showed strong bias for Gamma in moderate agreement with the modeling results (Table 3). HADDOCK predicted Gamma (159/200), Epsilon (25/200) and “Other” (15/200). The high number of Delta in RF2NA and Gamma in HADDOCK could potentially be better interpreted together in considering that these modes are minor variants of each other with equal positioning of the β1-β2-loop. The cross-platform model DockQ consistency comparisons for N.sub_nat_ (Fig 5) showed that a sub-set of the Epsilon models were virtually identical and reached high quality with excellent scores ≥ 0.8. The bulk of AF3 vs. Chai-1 model pairings, however, were in the medium quality range (DockQ ≈ 0.6), and all were in Epsilon, showing high cross-platform convergence on Epsilon. Both these platform models for N. sub_nat_ compared to RF2NA models had most pairings in the acceptable quality range and were consistently divergent on mode (i.e. “Different”) reflecting the high number of modeled Delta modes in RF2NA and Epsilon in the other two platforms.

#### B.den_nat_

All models by AF3 (165/165) and Chai-1 (91/91) were Epsilon, as were the majority (20/22) of the RF2NA models. In addition to Epsilon, RF2NA modeled a single Gamma (1/22) and a single “Other” (1/22) model. The AF3 models were significantly better in terms of PAE than those from Chai-1 whilst there was no significant difference in terms of pLDDT. Both Chai-1 and AF3 B.den_nat_ models were again significantly better than the RF2NA models. In the PAE/pLDDT scatter plot B.den_nat_ models overlapped with E.cor_nat_, N.sub_nat_, K.den_nat_ and N.muc_nat_ models in Epsilon and with E.cor_nat_, N.muc_nat_, and N.cin_nat_ in Gamma (Fig 4B and C). The re-docking in HADDOCK corroborated the modeling experiments in finding a strong bias for Epsilon (145/200), moderate number of Eta (“Other”) (54/200) and equivalently to RF2NA, only a single Gamma (1/200). The cross-platform model DockQ consistency comparisons for B.den_nat_ (Fig 5) showed that all AF3 vs. Chai-1 model pairings were in the medium DockQ quality range and convergently Epsilon, with some pairings being near high quality (Fig 5). All AF3 vs. RF2NA pairings were in the acceptable range, except for a residual of pairings that were in the lower medium range, divergent as Different or convergently in the Epsilon binding mode. All Chai-1 vs. RF2NA pairings were similarly also in the acceptable range with either Different or convergently Epsilon modes.

#### K.den_nat_

All models by AF3 (105/105) and Chai-1 (446/446) were Epsilon. AF3 models were significantly better than Chai-1 in regard of pLDDT and insignificantly different for PAE. RF2NA models were found in Zeta (69/98), Epsilon (25/98) and Gamma (4/98) with significantly poorer PAE/pLDDT quality scores than AF3 and Chai-1. In the PAE/pLDDT scatter plot a sub-fraction of AF3 and Chai-1 K.den_nat_ Epsilon models overlapped in addition to models of all other ComP_nat_s except N.cin_nat_ in Epsilon and all ComP_nat_s in Gamma (Figs 4B and C). Re-docking in HADDOCK predicted unambiguously Epsilon (200/200) in full agreement with AF3 and Chai-1 and moderately different from RF2NA. The high number of Zeta in RF2NA and Epsilon in HADDOCK could potentially be better interpreted together in considering in that these modes, like Gamma/Delta,are minor variants of each other with identical positioning of the β1-β2-loop. The cross-platform model DockQ consistency comparisons for K.den_nat_ (Fig 5) showed that some model pairings reached high DockQ ≥ 0.7 for AF3 vs. Chai-1 comparisons, but most pairings were in the acceptable range and convergently Epsilon. For the comparisons to RF2NA, most model pairings were in the acceptable range, yet some were in the medium range. The distributions of divergently Different and convergently Epsilon pairings were very similar.

#### N.muc_nat_

All models by AF3 (105/105) were Epsilon. No Chai-1 models reached ipTM ≥ 0.6, and all the 500 best scoring models were Epsilon (500/500). The relatively few successful RF2NA models were Gamma (10/11) and “Other” (1/11). The AF3 N.muc_nat_ models were significantly better than the Chai-1 models in terms of PAE and Chai-1 models were significantly better than AF3 in terms of pLDDT. The models from both Chai-1 and AF3 were significantly better than those from RF2NA in terms of PAE, but pLDDT was not significantly different between Chai-1 and RF2NA. In the PAE/pLDDT scatter plot N.muc_nat_ models overlapped with E.cor_nat_, N.sub_nat_, K.den_nat_ and B.den_nat_ models in Epsilon and all ComP_nat_ except N.sub_nat_ in Gamma (Fig 4B and C). Re-docking in HADDOCK corroborated findings from AF3 and RF2NA with a strong bias for Epsilon (145/200), yet both Gamma (53/200) and “Other” (2/200) modes were re-docked in concordance with RF2NA. The cross-platform model DockQ consistency comparisons for N.muc_nat_ (Fig 5) showed that some models reached high DockQ scores close to 0.8 for AF3 vs. Chai-1, with a broad range of DockQ scores, both acceptable and medium and convergently Epsilon. The comparisons to RF2NA were mostly in the acceptable range, represented by both divergent Different and convergent Epsilon modes.

#### E.cor_nat_

All models by AF3 (95/95) were uniquely Gamma. Chai-1 models were ambiguous in terms of binding mode in that the models with lowest ipTM were either Gamma (138/381) or “Other” (86/381), while the better models (157/381) were Epsilon. The RF2NA models were also ambiguous in terms of binding mode, with Gamma (26/81), Epsilon (23/81), Zeta (25/81), Delta (1/81) and Other (6/81) assignments making E.cor_nat_ the ComP_nat_ with most modeled modes overall (see Fig 3E). The Chai-1 models were significantly better than both AF3 and RF2NA and AF3 models were significantly better than RF2NA in terms of PAE/pLDDT. In the PAE/pLDDT scatter plot E.cor_nat_ models overlapped with N.sub_nat_, B.den_nat,_ N.muc_nat_ and K.den_nat_ models in Epsilon and with all ComP_nat_ except N.sub_nat_ in Gamma (Fig 4B and C). The re-docking in HADDOCK showed in stark contrast to AF3 and RF2NA no Gamma but a strong bias for Epsilon (170/188) and a few resembling Zeta (18/188). E.cor_nat_ was consequently the least coherently modeled ComP_nat_ for either Epsilon and Gamma across platforms and in re-docking. The cross-platform model DockQ consistency comparisons for E.cor_nat_ (Fig 5) showed that AF3 vs. Chai-1 paired with both medium and acceptable DockQ scores, most often convergently Gamma with some divergent and acceptable Different model pairings. In the AF3 vs. RF2NA comparisons (Fig 5), most pairings were in the acceptable range, convergently Gamma and divergently Different. The bulk of Chai-1 vs. RF2NA model pairs were in the acceptable range and convergent on Gamma, Epsilon and Other modes. Also, several divergently Different comparisons were found.

#### N.cin_nat_

The AF3 models never reached ipTM ≥ 0.6 and of the 100 best scoring models, were Gamma (96/100) and few Epsilon (4/100). In Chai-1, a unique Theta mode was observed for all models (395/395), a mode exclusively seen in Chai-1 for this N.cin_nat_. The RF2NA models were either Gamma (40/52) or the resembling Delta (12/52). Chai-1 models were significantly better than AF3 and RF2NA in terms of PAE and pLDDT. AF3 models were significantly better than RF2NA in terms of PAE while interestingly, RF2NA models were significantly better than AF3 models in terms of pLDDT. In the PAE/pLDDT scatter plot N.cin_nat_ overlapped with N.men_nat_ in Epsilon and all ComP_nat_s except N.sub_nat_ in Gamma (Fig 4 B and C). In the unique Theta models, a complete strand separation in one end of the DNA and one flipped out base were observed (Fig 3 F and Fig S1 C), again showcasing a unique Chai-1 feature for altering DNA structure, possibly to induce an optimal fit. Re-docking in HADDOCK showed like AF3 and RF2NA a bias for Gamma (121/200) with less Epsilon (79/200). Ambiguities in both modeling and re-docking and the special strand-splitting Theta mode predicted by Chai-1 suggests that N.cin_nat_ has unique features in its structure to significantly affect both modeling and re-docking of the AT-DUS template. The cross-platform model DockQ consistency comparisons for N.cin_nat_ (Fig 5) showed that AF3 vs. Chai-1 comparisons never reached acceptable DockQ, and all pairings were divergently Different. Several AF3 vs. RF2NA pairings reached high-quality DockQ, albeit with most pairings in the medium range and converged on Gamma. Several models were divergently in Different modes. All Chai-1 vs. RF2NA comparisons were below the acceptable range and consistently divergent with respect to mode.

#### N.men_nat_

The AF3 models never reached ipTM ≥ 0.6, and the 100 best scoring models were Gamma (76/100), Epsilon (21/100) and Delta (3/100). The relatively few Chai-1 models were in Epsilon (12/13) and Zeta (1/13) and the RF2NA models were in Gamma (56/91) and Delta (35/91). Chai-1 gave significantly better PAE and pLDDT scores than both AF3 and RF2NA. As with Ncin_nat_, the RF2NA models were significantly better than the AF3 in terms of pLDDT, albeit having significantly weaker PAE. In the PAE/pLDDT scatter plot N.men_nat_ models overlapped with N.sub_nat_, K.den_nat_ and N.cin_nat_ models in Epsilon and K.den_nat_, N.cin_nat_, E.cor_nat_ and N.muc_nat_ in Gamma (Fig 4 B and C). Re-docking in HADDOCK showed in contrast to AF3 and RF2NA a bias for Epsilon (130/200), yet with several Gamma (68/200) and a few Zeta (2/200). Few successful Chai-1 models may signify the platforms difficulty in modeling this ComP_nat_ complex. The cross-platform model DockQ consistency comparisons for N.men_nat_ (Fig 5) showed that most AF3 vs. Chai-1 pairings were in the acceptable range, being Different or convergently Epsilon, with some Epsilon pairings clustering in the medium range. AF3 vs. RF2NA convergently predicted Delta and Gamma in the acceptable and medium ranges with some pairings close to the high-quality range. All Chai-1 vs. RF2NA pairings were divergently Different and in the acceptable DockQ score range.

### AF3 modeling quality in matching native vs. scrambled DUS to ComP

In order to explore modeling quality and consistency in regard of DUS specificity, we modeled ComP_nat_s and ComP_scr_ and compared the resulting ipTM and Chain Pair PAE minimum (CPPM) distributions for all models (Fig 6 A and B; Code output S2).

**Figure 6.**
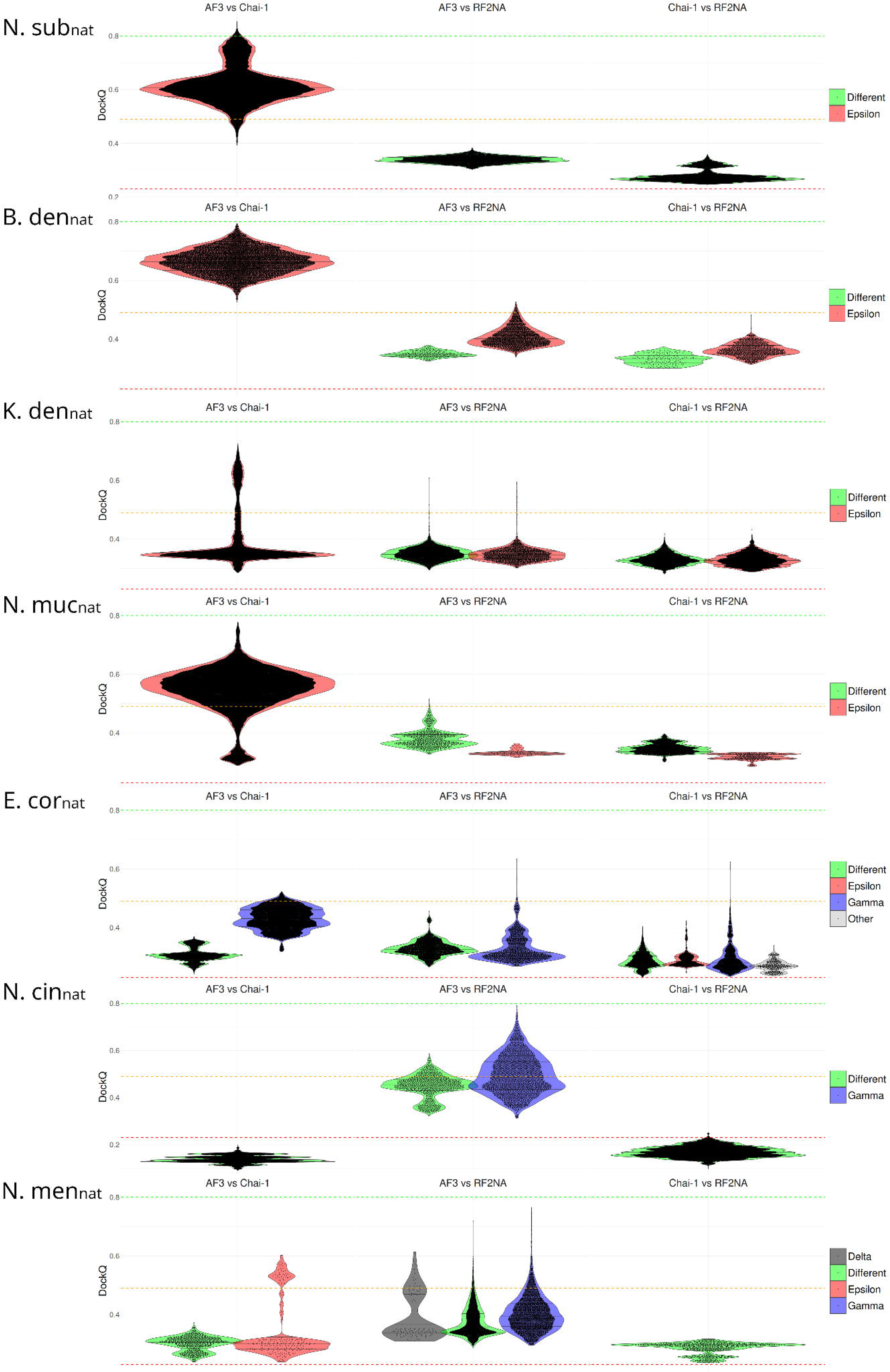
A: Violin plots showing distributions of ipTM for native (“Native”) and scrambled DUS (“Scrambled”) runs in AF3 for the docking set consisting of five DUS dialects encompassing seven ComP_nat/scr_ models. Box plots encompassing the median and interquartile ranges are seen inside the violins. AA-king3DUS (K.den). AG-DUS (B.den + N.sub). AG-eikDUS (E.cor). AG-mucDUS (N.muc). AT-DUS (N.men + N.cin). **B:** Same as in A for CPPM. Refer to Code Output S2 for statistical significance tests for Native vs. Scrambled for all DUS dialects.

Since the native DUS dialects were the same in the two pairs N.men_nat_/N.cin_nat_ (AT-DUS) and B.den_nat_/N.sub_nat_ (AG-DUS), these were combined in the assessment. AF3 was chosen as platform as it covered most ComP_nat_ (6/7) with robust (ipTM ≥ 0.6) models. Wilcoxon rank sum tests (Code output S3), using significance level α=0.05, showed that the ipTM distributions (Fig 6 A) for native DUS models (“Native”) and scrambled DUS models (“Scrambled”) were significantly different in all comparisons except for AA-king3DUS. Furthermore, ipTM in Native was significantly better (higher) than Scrambled for AG-DUS,

AG-eikDUS and AG-mucDUS. In contrast, for AT-DUS the ipTM distribution was significantly lower (inferior) in Native than in Scrambled. This suggested a greater difficulty for AF3 to generate high-quality models for Scrambled than Native for AG-DUS, AG-eikDUS and AG-mucDUS, whereas AF3 modeled high quality models for both Scrambled and Native for AA-king3DUS. For AT-DUS AF3 modeled better complexes with Scrambled than with Native. The CPPM scores (Fig 5 B) reflected ipTM in that the distributions were significantly different in all comparisons except AA-king3DUS. CPPM was significantly better (lower) in Native than Scrambled for AG-DUS, AG-eikDUS and AG-mucDUS. For AT-DUS, CPPM was significantly higher (inferior) in Native than Scrambled, corroborating the results for ipTM and reflecting AF3’s difficulty in predicting ComP_nat_s with the AT-DUS dialect and high variation in the models for these complexes. The CPPM results also indicated that AF3 was not distinguishing AA-king3DUS from its scrambled versions. The frequencies of assigned binding modes using scrambled DUS in AF3 are shown in Table 4, showing that also these are mainly Epsilon and Gamma.

**Table 4.**
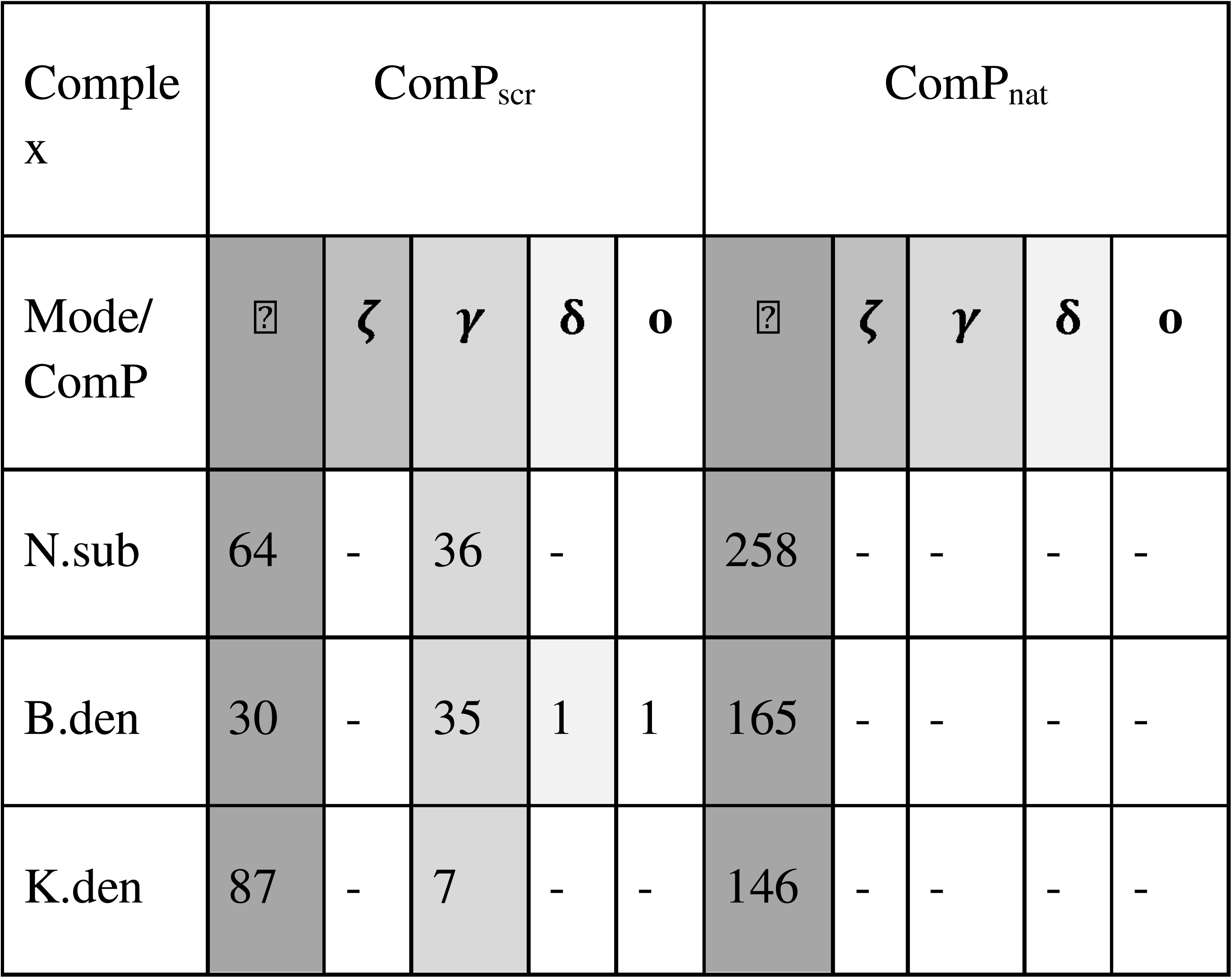
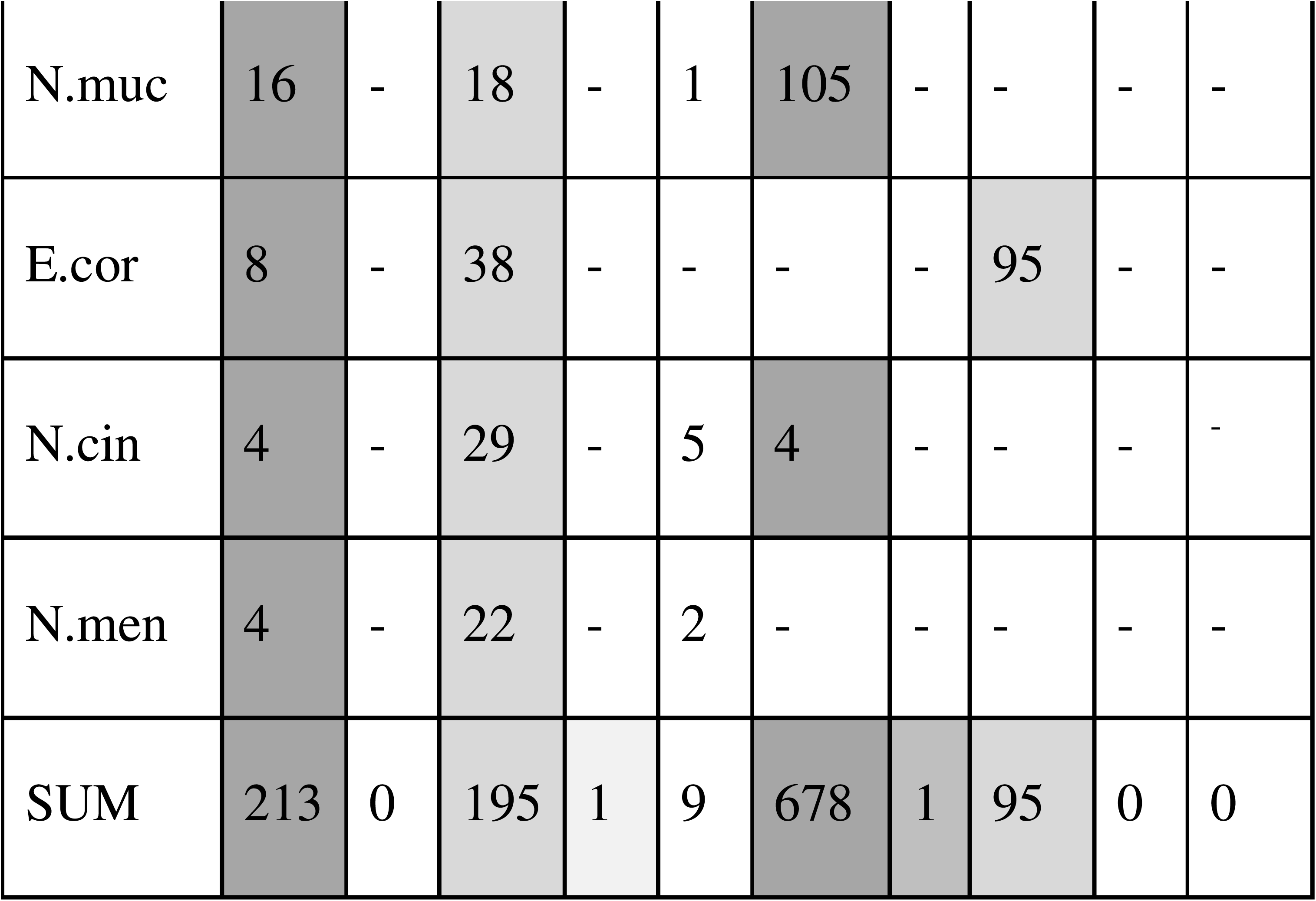
Binding mode distribution across all seven ComPs generated with either scrambled DUS (left) and native DUS (right) in AF3. Greek letter symbols for modes are ⍰ = Epsilon, ***ζ***= Zeta, ***γ*** =Gamma, ***δ***=Delta and **o**=other. Data for native DUS are as in Table 1 but with a strict ipTM ≥ 0.6 applied.

In considering the mode differences for each ComP going from native to scrambled DUS, as described in Table 4, we observe the following tendencies: Robust Gamma models appear in six ComP_scr_ (N.sub_scr_, Kden_scr_, Bden_scr_, N.muc_scr_, N.cin_scr_, N.men_scr_) where there were no Gamma modes above cut-off for ComP_nat_, and inversely Epsilon appear in E.cor_scr_ where there were none in E.cor_nat_. AF3 generated robust models for ComP_N.men_ only when using scrambled DUS and below cut-off with native DUS. Representative ComP_scr_ PAE-plots for each mode and ComP_scr_ are shown in Fig S7.

## Discussion

In this study, three different deep learning protein structure prediction platforms and one traditional molecular docking platform were used to model the interaction between ComP and DUS in native ComP_nat_ complexes. The seven ComPs studied were chosen as they represented both a diverse set regarding primary structure from three different genera (*Neisseria/Bergeriella, Eikenella* and *Kingella*) (four genera if considering *Bergeriella* distinct from *Neisseria*) and had their respective genomes enriched with five different DUS dialects as previously described (Frye, 2016). The overall 3D structure of the ComP protein was found conserved in these investigated ComPs. The overall ComP domain organization and structure conservation were found highly similar and showed that all tested proteins likely were functionally conserved as DUS specific receptors as previously interpreted from ComP sequence alignments (Cehovin et al., 2013). Differences in amino acid composition of ComPs, as well as minor differences in structural domains in ComPs, e.g. the here predicted α-turns in AF3 ComP_B.den_, ComP_E.cor_, and ComP_N.muc_, might together help understand general and specific characteristics of the ComP :: DUS interaction. It remains unclear why RF2NA generally did not predict the highly conserved disulfide bridge between β1-sheet and the DD-region. The disulfide bridged DD-region is of particular interest as it is expected to facilitate specific DUS-binding (Cehovin et al., 2013). Although the disulfide bridge between the β1-sheet and the DD-region is missing in RF2NA models, the relative positioning of the DD-loop in the overall ComP structure remains largely the same in the RF2NA models as in all other modeled structures and seems therefore not to have had significant impact on the RF2NA ComP_nat_ models. It cannot be fully excluded however, that the propensity to model individual DNA binding modes by RF2NA could be influenced by more potential flexibility of the DD-loop lacking its disulfide bridge.

### The ComP_nat_ binding modes

By modeling ComP_nat_s, six different binding modes and one heterogeneous “Other” category were described. These categories were based on their primary and main interactions of two consistently DNA-interacting loops in ComP, the α1-β1 loop and the β1-β2 loop. Epsilon/Zeta/Theta were categorized by how these two loops interact through the DNA major groove and Gamma/Delta through the minor groove (Table 2). The sub-divisions of Epsilon and Gamma into Zeta/Theta and Delta, respectively, were based on how the DD-region differentially interacted with DNA. Although ComP :: DUS docking by HADDOCK in our hands frequently distorted the DNA structure making the grooves challenging to distinguish, we have interpreted previously illustrated results from in silico ComP-DUS docking experiments in Berry et al. (2016) and Hughes-Games (2020) to reflect the Epsilon mode, i.e. main interactions through the major groove. Hence the Epsilon mode as such, has some previous docking support.

Strong support for the Epsilon mode was also found here across platforms and for different ComP_nat_. All ComP_nat_ were modeled in Epsilon by one or more platforms. Epsilon was the most frequently modeled mode in AF3 and Chai-1 by a considerable margin, and this therefore supports Epsilon as a very likely ComP_nat_ complex. All platforms modeled Epsilon for three or more ComP_nat_ and the Epsilon models were particularly robust and consistent in AF3 and Chai-1 for N.sub_nat_, N. muc_nat_, B. den_nat_ and some of the K. den_nat_ (Table 3; Fig3-5). Fewer successful Epsilon models were found in E. cor_nat_, N. cin_nat_ and N. men_nat_, for which more Gamma was observed.

Zeta resembles Epsilon by having the α1-β1 and β1-β2 loops entering the major groove. Zeta was not found in AF3. Zeta was always modeled together with Epsilon for the same ComP_nat_ in the two platforms where Zeta was seen, Chai-1 and RF2NA, signifying the similarity of the two modes. A single Zeta model was found for N. men_nat_ in Chai-1 together with several Epsilon. In RF2NA, Zeta and Epsilon were modeled similarly for K. den_nat_ and E. cor_nat_. These observations support considering Zeta a DD-loop variant of Epsilon.

The Theta mode was uniquely identified in N.cin_nat_ and only in Chai-1. Unlike all other modes, Theta showed major conformational changes in the DNA, complete strand separation, which potentially could allow better protein access to DNA bases. Theta resembles Epsilon/Zeta in that the major interactions were in the major groove, which was pried apart in Theta. The consistently flipped-out base is the central thymine in the conserved 5’-CTG-3’ DUS motif, alluding to involvement in DUS-specific binding. The only other severe alteration of the DNA structure was found in N. sub_nat_ Epsilon Chai-1 models, also displaying a flipped-out base. This fifth thymine base immediately precedes the conserved CTG core of the AG-DUS and it is thus tempting to speculate that the modeled conformational change in the DNA may instigate specific binding. The capability of Chai-1 to model substantial conformational changes in DNA sets it apart from AF3 and RF2NA and may be due to different emphasis on confidence metrics and that Chai-1 allowed also defining different MSA schemes for the DNA. Further experimentation by e.g. molecular dynamics is required to investigate these models in depth by manipulation of input parameters and MSA schemes.

Particularly strong support for Gamma was found in three ComP_nat_, N. cin_nat_, N. men_nat_ and E. cor_nat_. Together all platforms modeled Gamma for at least one of these three ComP_nat_ and all ComP_nat_s were modeled in Gamma in at least one platform. RF2NA generated Gamma models for all ComP_nat_s. Gamma was the most frequently modeled mode in RF2NA and second most frequent mode in AF3 and Chai-1 after Epsilon.

Like Zeta/Theta can be considered variants of Epsilon, Delta resembles Gamma in having its main interactions with the minor groove. When Delta was found for any ComP_nat_, the same platform also modeled Gamma, showing a correlation between the two modes as with Epsilon/Zeta. These mode and sub-mode models may reflect a flexibility of the DD-region which may be relevant, particularly in regard to the missing disulfide bridge in RF2NA. Delta was modeled for N.sub_nat,_ N.cin_nat,_ N.men_nat_ and E.cor_nat_ by RF2NA and below ipTM cutoff in AF3 (N.men_nat_). N.sub_nat_ was the only modeled complex in this RF2NA Delta subset with also very robust Epsilon models in AF3 and Chai-1. In modeling predictions yielding Delta, N.cin_nat,_ N.men_nat_ and E.cor_nat_ stand out as complexes imposing modeling challenges for all platforms, although E.cor_nat_ achieved moderately high ipTM scores in Gamma in AF3 and Gamma/Epsilon in Chai-1. E. cor_nat_ was also the only complex with a substantial fraction uncategorized modes assigned Other.

The Eta model was never modeled but appeared in the re-docking experiments using HADDOCK. Eta was therefore not considered a supported mode but draws attention to the ability of re-docking to both reproduce other modeled complexes and predict new. HADDOCK did robustly generate re-docks showing both Epsilon and Gamma modes in support of these.

Other modes than the above were considered anomalies in modeling. As can be inspected in Fig 3, the Other modes of E.cor_nat_ in Chai-1 and a single model of B. den_nat_ in RF2NA either bind DNA to the opposite end of ComP or at greater distance and tilt relative to all other models. If anything, these models again show the challenge in convergently modeling E.cor_nat_ to any mode (modeled in all modes except Theta) and that the RF2NA outputs generally are more mode-variable than AF3 and Chai-1.

### Platform performances and consensus predictions

Generally, under our application of confidence scores and metrices, RF2NA predicted less robust models than AF3 and Chai-1. This observation aligns with those of Abramson et al. (2024) demonstrating RF2NA to be inferior to AF3 for protein-nucleic acid prediction on benchmark sets of the PDB database, having about 40% lower success rate than AF3. However, since the ComP :: DUS structure remains unresolved and may be unusual due to its unique extracellular confinement, it cannot be excluded that RF2NA models may be informative for identifying the DUS-specific mode. Abramson et al. (2024) does not describe if there was a perfect overlap in successful AF3 and RF2NA models or if RF2NA modelled certain complexes or types of complexes with greater success than AF3. However, the systematically high PAE values of successful RF2NA models contrasts the unity of nearly identical AF3 models with much lower PAE values and speaks at least for the robustness of the AF3 models and possibly the impropriety of emphasizing PAE metrics for RF2NA models. In support of the RF2NA Gamma models, it is notable that the pLDDT scores from RF2NA generally were relatively high (>82), like most of the models from AF3 and Chai-1 and that N.cin_nat_ and N.men_nat_ modeled with significant better pLDDT values than AF3 in Gamma. Although there is some variation within successful Chai-1 models, the consistent overlap with AF3 in terms of both Epsilon and Gamma is evident. Higher variation in the Chai-1 models compared to those from AF3 may also be explained by Chai-1’s variable MSA input, whereas the AF3 web server does not allow for MSA customization (AF3 now released as open source). Thus, the cross-platform deviance may stem from the three platforms’ unequal algorithmic modeling, score metric emphases and source inputs or simply that AF3 and Chai-1 models being of generally better quality than the those from RF2NA. However, all platforms and modes may prove informative when exploring DUS-specificity. Assuming that there is one and only one DUS-specific mode across all ComPs, it is paradoxical that even in the cases where the models were of putatively high quality in both programs (RF2NA: PAE < 10; AF3: ipTM ≥ 0.6), the modes were robustly either Epsilon or Gamma. For instance, RF2NA may have provided insight into ComP_nat_s modeled with AT-DUS (N.men_nat_ and N.cin_nat_) with solely Gamma/Delta predictions, as both of these had models of PAE <10, while AF3 always failed to predict N.men_nat_ and N.cin_nat_ models above its ipTM threshold. At the same time, we never observed any RF2NA models of higher quality than any AF3 or Chai-1 models when considering confidence from PAE, only pLDDT cases and DockQ consistency, which is striking. RF2NA predicted generally more modes per ComP_nat_ than AF3 and Chai-1. RF2NA therefore had challenges with converging to individual modes and displayed greater modelling flexibility relative to AF3 and Chai-1. This may be due to fundamentally different emphasis on different confidence metrices. Although more modes were modelled per ComP_nat_ in RF2NA, a converging trend towards Gamma (and Delta) was observed in support of a robust loop-interaction prediction with the minor groove. Although it is unclear to which degree RF2NA with its modelling flexibility can return “false positive” ComP_nat_ models of high confidence that fail to recover the true binding mode, Baek et al. (2024) reported that 81% of models with PAE <10 correctly predicted the interface of the complex, which is substantial.

In exploring the AF3 modeling quality of DUS against scrambled DUS using CPPM confidence, ComP_N.cin_ and ComP_N.men_ were found as the only ComPs where scrambled DUS models were significantly better than their native pairings of which most were below the ipTM threshold (Table 3-4; Fig 5). Most of these scrambled models were in Gamma (Table 4). For AF3 to predict robust models for these two ComPs, a scrambled AT-DUS was required, suggesting that individual bases in the native AT-DUS were preventive for AF3. In contrast, ComP_B.den_/ComP_N.sub_, ComP_N.muc_ and ComP_E.cor_ AF3 models were significantly better for native Epsilon or Gamma complexes than scrambled. These results suggest that

AF3 more accurately predicted the DUS-specific mode than any non-specific mode in these ComPs although they were discrepantly Epsilon and Gamma and hence not parsimonious in regard of mode. The ComP_scr_ data show that AF3 predicted Gamma for N.men_scr_ and N.cin_scr_ in a seemingly DUS-independent manner and E.cor_nat_ in a seemingly DUS-dependent manner (Table 4). In contrast, AF3 robustly predicted Epsilon for B.den_nat_, K.den_nat_, N.sub_nat_, N.muc_nat_ in a seemingly DUS-dependent manner and more Gamma in a seemingly DUS-independent manner. Furthermore, ComP_K.den_ did not significantly differentiate between its native DUS AA-king3-DUS and the scrambled versions (Fig 5), questioning whether any of the two modes could represent the ComP_K.den_ DUS-specific mode. We have previously shown that the DUS dialect phylogenetic distribution is consistent with the robust core genome Neisseriaceae phylogeny (Frye, 2013). Finding here that the seven different ComP_nat_s showed propensities to model the two different modes in a manner inconsistent with the ComP phylogeny remains puzzling. The strict conservation of the essential CTG core of DUS suggests the molecular interactions and mechanism responsible for the ComP :: DUS binding and consequential DNA uptake is one. Wet-lab experiments have shown that DNA containing any single mutation in this CTG core completely and uniquely abrogates transformation in both *N. meningitidis* and *N. subflava* (Berry et al., 2013; Frye et al., 2013) whose ComPs here show propensities to model either Epsilon (*N. subflava*) or Gamma (*N. meningitidis*) with their respective DUS in AF3 and other platforms. These *in vivo* results support the notion that there is only one DUS-specific interaction and mechanism. The modeling platforms used here do either inconsistently or not at all capture this expected parsimony in their robustly predicted different modes. There could be different reasons for this. Firstly, all predicted models are precisely that, predicted models. Although the here applied platforms represent state-of-the-art tools for protein structure and protein-DNA complex structure prediction, obtaining functional understanding is not immediate or straight forward (Varadi et al., 2025). Issues with confidences of predicted disordered protein regions, dynamic interactions, multi-chain complexes such as protein-DNA, training-set representativeness and lack of benchmarking studies are different recently discussed challenges (Varadi et al., 2025; Chakravarty et al., 2024; Ruff & Pappu, 2021). The ComPs investigated here are to platform variable extents modeled based on pattern recognition, learned protein energetics and possibly wet-lab resolved 3D structures inside user-inextricable black-box algorithms. Basing our analysis on different such complex deep-learning platform algorithms with diverging dependencies and potentially dynamic inputs, we cannot exclude the possibility that the positioning of particularly flexible or disordered regions in ComP are inaccurately modeled, such as the lack of biologically important disulfide bridges in RF2NA. We also observe varying abilities to severely modulate the structure of DNA, particularly seen in some Chai-1 models. A possible consequence of any such platform biases could be that one mode is artificially favored in modeling not to represent an expected ground truth. Hence, our search for cross-platform consensuses applying fundamentally different modeling algorithms. As shown in Fig 2 the topologies of the investigated ComPs are highly similar, yet three ComPs have modeled α-turns different from the others. The β1-β2 loop have overlapping α-turns in ComP_Ecor_, and ComP_Nmuc_ and ComP_Bden_ have one in the DD-loop. We cannot find that these α-turn divergent topologies could explain propensities for either Epsilon (ComP_Nmuc_ and ComP_Bden_) or Gamma (ComP_Ecor_) in any systematic way and we have fair confidence in their predicted protein 3D structures and overall structural coherence across platforms.

All our predicted models and modes are static snapshots which do not reflect any spatiotemporal dynamics of the biological ComP :: DUS interaction. We cannot therefore exclude the possibility that the DNA binding modes identified here may be limited to showing the initial interactions between DUS and ComP, i.e. the first step towards making an interaction which involves the essential CTG-core after sequential or simultaneous conformational changes have been made in the protein, DNA, or both. We do note that it is particularly the two AT-DUS binding ComPs and the AG-eikDUS binding ComP which are more difficult to model in Epsilon than Gamma, particularly in AF3. The AT-DUS differs from AG-DUS only in position −1 (A<u>T</u>GCCGT<u>CTG</u>AA vs. A<u>G</u>GCCGT<u>CTG</u>AA) five nucleotide positions away from the CTG core which occupy DUS-positions 6-8. Hence the singular −1 position away from the conserved core seems restrictive for making a robust Epsilon mode in AT-DUS ComPs and should be investigated further (see below). The AG-eikDUS (AGG<u>CT</u>AC<u>CTG</u>AA) dialect differs in 2-3 positions relative to the other dialects and two of these differences are dialect-unique with T in position 3 and A in position 4 much closer to the CTG-core than −1. For ComP_Nmen_, Epsilon modes appear in AF3 only when scrambled AT-DUS are used and relatively few Epsilon modes are found for ComP_Ncin_ with both templates (Table 4). ComP_Ecor_ with AG-eikDUS and scrambled versions show similarly that Epsilon appears only when using scrambled AG-eikDUS in AF3. Consequently, all three ComPs harbour the propensity to model Epsilon *per se*, but are restricted by their native DUS to do so. It is interesting that Epsilon modes remain scarce relative to Gamma in models of all these three ComPs when using scrambled versions of DUS. This shows that these ComP’s retain an overall propensity for Gamma with different DNA templates even though their DUS dialect-unique positions (−1 and 3+4) are not the same. The other four ComPs (ComP_Kden_, ComP_Bden_, ComP_Nsub_ and ComP_Nmuc_) show an opposite tendency to loose their strict Epsilon propensity when scrambled DUS versions are used and Gamma modes appear (Table 4). These Epsilon to Gamma mode-shifts are particularly strong for ComP_Nmuc_ and ComP_Bden_ which show about equal numbers of Gamma and Epsilon modes when scrambled DNA templates are used, and are phylogenetically closer to ComP_Nmen_ and ComP_Ncin_ than ComP_Kden_ and ComPE_cor_. In contrast, ComP_Kden_, retain much of its Epsilon propensity when the scrambled AA-king3DUS was used resembling the disinclination to shift from Gamma to Epsilon as discussed above. Since Epsilon and Gamma models are robustly predicted with scrambled DUS in all investigated ComPs we find support for the notion that the models predicted here could reflect initial contacts where DUS-specific interactions are yet to completely unfold. Further systematic modeling studies of propensities to model different modes using non-native DUS, artificial deconstructed DUS (incl. all 36 possible individual transitions and transversions pr. DUS dialects) and DNA templates of different lengths and internal DUS-positions could shed further light on these shifts in mode propensities and better support Gamma or Epsilon as the one parsimonious DNA-binding mode responsible for DUS-specificity. Any mode further supported by in silico approaches would require confirming wet-lab assaying experiments to which we here provide direction.

Taken together, these data support that Gamma and Epsilon reflect the most robust and inherent DNA-binding properties of ComPs irrespective of DUS-specificity. Experimental data by Cehovin et al. (2013) showed that, in addition to DUS-specific transformation, non-DUS transformation is also mediated by ComP_N.men_. These *in vivo* results showed not only the well-characterized wild-type bias for DUS-specific transformation, but also that non-DUS transformation could reach the level of DUS-specific transformation when ComP was overexpressed (Cehovin et al., 2013). We hypothesize that this DUS/non-DUS ambiguity could be connected to the two distinct and robust binding modes identified here, Epsilon and Gamma. Finally, DNA binding and DNA uptake are consecutive events in the transformation process of which the modeling performed here considers the first step and initial interaction between protein and DNA. Specific DNA-binding may require considerable conformational changes in either or both ComP and DUS of which Gamma/Delta and Epsilon/Theta/Zeta, could represent initial states. Molecular dynamics simulations may shed further light on downstream conformational changes in binding partners in future in silico experiments and ultimately resolve the specific molecular interactions responsible for DUS-specificity and better explain the here documented divergent modeling performance of the different platforms.

## Summary of conclusions

Two distinctly different DNA binding modes of ComP, Epsilon and Gamma, were robustly predicted by AF3, Chai-1 and RF2NA and HADDOCK. Epsilon and Gamma differ characteristically in how two conserved ComP protein domains, the β1-β2-loop and the DD-loop, interact with the minor and major grooves in the DNA. Although robustly modeled across platforms, the modeling platforms showed considerable variation in their propensity to model these two modes in the seven ComP :: DUS complexes explored here. RF2NA seemed to place fundamentally different emphasis on PAE relative to AF3 and Chai-1 which skewed the quality assessment in a systematic manner to the disadvantage of RF2NA models. In contrast to PAE, pLDDT scores were considerably more overlapping and within high confidence range (>82) across platforms in Epsilon and Gamma models and notably so in Gamma. Robust support for Epsilon was found in N. sub_nat_, B. den_nat_, K. den_nat_ and N. muc_nat_, and Gamma mainly found its support in E. cor_nat_, N. cin_nat_ and N. men_nat_. RF2NA generally predicted more Gamma whilst AF3, Chai-1 and re-docking in HADDOCK predicted more Epsilon to show inherent differences between the three modeling platforms and relative to re-docking. No parsimonious Epsilon vs. Gamma mode distribution could be identified to elucidate if any of these modes could represent DUS-specific or DUS-independent binding. AF3 predicted robustly both Epsilon and Gamma models of higher quality (CPPM) when using native DUS relative to scrambled DUS and possible modeling limitations responsible for these mode-incongruent results are discussed. Scrambling the DUS provided models shifting their native propensities to bind either Epsilon or Gamma to the other mode, showing inherent modeling capabilities of all ComPs to accommodate both modes irrespective of DUS-specificity. Although six different binding modes were predicted here, we find only strong consensus modeling support for all ComPs to bind DNA in Epsilon and Gamma.

## Data availability

Scripts and data used are provided in the GitHub repository https://github.com/stianale/ComP-DUS-modeling.

## Author contributions

SAH: Data curation, formal analysis, investigation, methodology, implementation of computer codes and supporting algorithms, data presentation, writing original draft and review & editing. AHL; KA and SAF: Supervision of SH and scientific critical review, writing – review & editing. OHA: Conceptualization, funding acquisition, project administration, resources, supervision, visualization, writing – review & editing. A Ph.D. grant was provided for SAH from Faculty of Health Sciences, OsloMet, Norway. The funders had no role in study design, data collection and analysis, decision to publish, or preparation of the manuscript.

## Supporting information captions

**Table S1** ComPs used for the different species and the paired DUS-dialect.

**Table S2** The different MSA schemes used in Chai-1, listing the various settings.

**Figure S1** Representative ComP-DUS structures showing six different binding modes Epsilon, Gamma, Theta, Zeta, Delta and Eta modeled in AF3, Chai-1 and RF2NA.

**Figure S2** Alignment of the ComPs, using default settings in MAFFT v. 7.505, visualized in MVIEW v. 1.68. First 28 amino acids of the N. meningitidis mature ComP are trimmed and the other ComPs accordingly.

**Figure S3** Selected high-ranking HADDOCK models, representative of the most abundant binding mode of each ComP_nat_. A: N. muc_nat_ (Epsilon). B: N. men_nat_ (Epsilon). C: K. den_nat_ (Epsilon). D: E. cor_nat_ (Epsilon). E: B. den_nat_ (Epsilon). F: N. cin_nat_ (Gamma). G: N. sub_nat_ (Gamma). AF3, Chai-1 and RF2NA DNA binding modes were used to guide the assignment of grooves in the HADDOCK distorted DNAs.

**Figure S4** Representative PAE plots of the top ranking native AF3 model for each ComP_nat_ and assigned mode.

**Figure S5** Representative PAE plots of the top ranking native Chai-1 model for each ComP_nat_ and assigned mode.

**Figure S6** Representative PAE plots of the top ranking native RF2NA model for each ComP_nat_ and assigned mode.

**Figure S7** Representative PAE plots of the top ranking scrambled AF3 model for each ComP_scr_ and assigned mode.

**Code Output S1** Wilcoxon rank-sum tests on PAE and pLDDT, comparing AF3, Chai-1 and RF2NA. Median PAE and pLDDT values are also shown.

**Code Output S2** Wilcoxon rank-sum tests on ipTM and CPPM for each ComP_nat_.

**Code Output S3** Wilcoxon rank-sum test on DockQ scores for the internal platform consistency check.

