## Supporting information for "Two different and robustly modeled DNA binding modes of Competence Protein ComP - systematic modeling with AlphaFold 3, RoseTTAFold2NA, Chai-1 and re-docking in HADDOCK"

**Table S1** ComPs used for the different species and the paired DUS-dialect.

| Species | GenBank/RefSeq accession | ComP primary structure (N-terminally truncated) | DUS-dialect | DUS-dialect name |
|---|---|---|---|---|
| Neisseria subflava NJ9703 | EFC51080.1 | RSANLRAAHAALLENARFMEQFYAKKGSFKLTSTKW PELPVKEAGGFCIRMSGQAKGILEGKFTLKAVALDRE AEPRVLRLNESLTAVVCGKMKGKGSCTDGEEIFRGN DAECRPFTG | AGGCCGTCTGAA | AG-DUS |
| Bergeriella denitrificans NCTC10295 | STZ76941.1 | REGRLREAQAALLENAQFLEKHYRQTGSIRANSTTW PTLPVTEAGGFCIRLSGLARGQSNQTEGKFTLKAVAL DKTREPRVLKTNEALMTTICESSSSSCDDGLQHFSG DGSTDQDCRVYQH | AGGCCGTCTGAA | AG-DUS |
| Neisseria mucosa ATCC 25996 | EFC87706.1 | RDSEMRQALAALVESAQFMERFYQQNGSFKKTSTA WPDLPNSRSLENFCIYPHGLARGALDGKFTLKAVALD KNKEPRVIKINESLTTFICESTASSCDDVTKNYFSGAD KNCSVYRL | AGGCCGTCTGAA | AG-mucDUS |
| Eikenella corrodens ATCC 23834 | EEG23425.1 | RKSRLEEANAALLENSRAMERFYARNRTFKATSTTW PALAVSQTQHFCIKFQGNARGVLGDKYTIKAVAFDVS KEPRVLLINQDQTVRICQSSRSRCDNKEVFSGGNNID QECELLH | AGGCTACCTGAA | AG-eikDUS |
| Kingella denitrificans ATCC 33394 | EGC16987.1 | RKSRLSEVQQLMLDNAQAWERHYAAHGHYRQTSRK WAALPVQGNDDFCIRPQGAPRGAAHDGQYSLKAVA LDKTKEPRVLVMNQDLTFLLCEESSSTCAETDYFANP ARADKNCRSYP | AAGCAGCCTGCA | AA-king3DUS |

| | | | | |
|---|---|---|---|---|
| Neisseria meningitidis MC58 | WP_002214937.1 | EKAKINAVRAALLENAHFMEKFYLQNGRFKQTSTKWPSLPIKEAEGFCIRLNGIARGALDSKFMLKAVAIDKDKNPFIIKMNENLVTFICKKSASSCSDGLDYFKGNDKDCKLLK | ATGCCGTCTGAA | AT-DUS |
| Neisseria cinerea ATCC 14685 | WP_003678313.1 | EKARISAVRSALLENAHFMEKFYLQNGTFKQTSTKWPKLPIQEAEGFCIRLNGVARGALDSKFMLKAVAIDKNKEPRIIKMNENLVTFVCKGSTSSCDDGLDYFRGNDKGCTLFK | ATGCCGTCTGAA | AT-DUS |

**Table S2** The different MSA schemes used in Chai-1, listing the various settings.

| MSA scheme | MSA for ComP | MSA for DUS | Origin of MSA for ComP | Origin of MSA for DUS | Paired MSA | ComP MSA query coverage |
|---|---|---|---|---|---|---|
| 1 | Yes | Yes | BFD | Genomic DUS counts | Yes | 50.00% |
| 2 | Yes | Yes | Uniref30 | Genomic DUS counts | Yes | 50.00% |
| 3 | Yes | Yes | BFD | Genomic DUS counts | Yes | 75.00% |
| 4 | Yes | Yes | Uniref30 | Genomic DUS counts | Yes | 75.00% |
| 5 | Yes | Yes | BFD-Uniref30 composite | Genomic DUS counts | Yes | 75.00% |
| 6 | Yes | No | BFD | NA | No | 50.00% |
| 7 | Yes | No | Uniref30 | NA | No | 50.00% |
| 8 | Yes | No | BFD | NA | No | 75.00% |
| 9 | Yes | No | Uniref30 | NA | No | 75.00% |
| 10 | Yes | No | BFD-Uniref30 composite | NA | No | 75.00% |

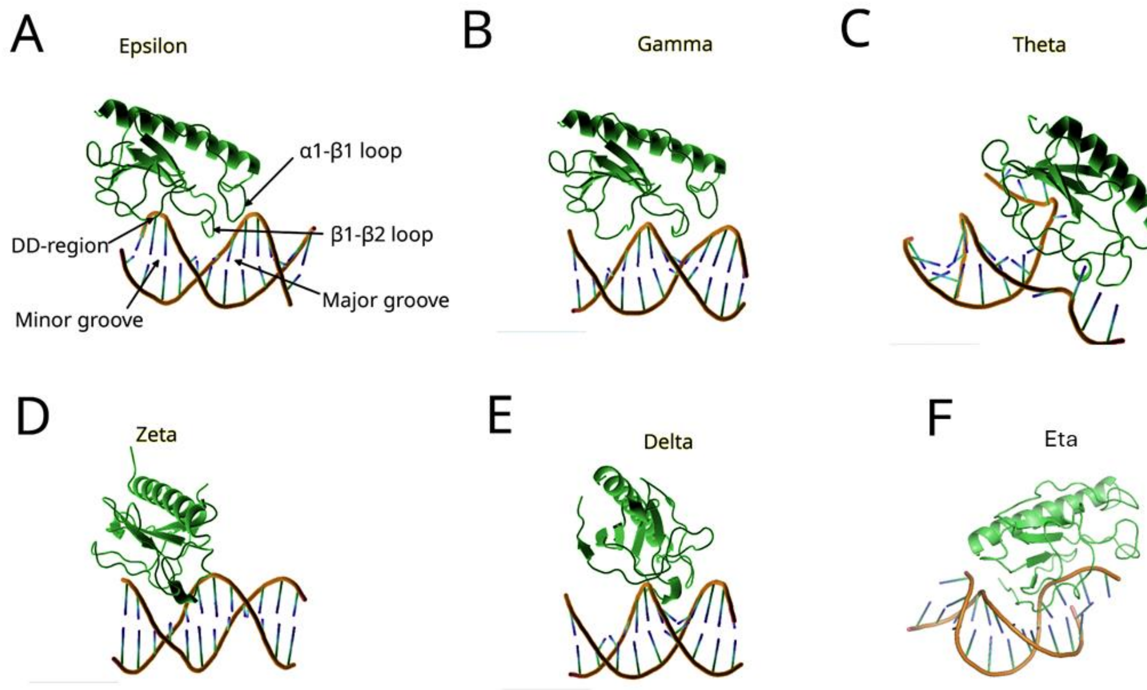


**Figure S1.** Representative ComP-DUS structures showing six different binding modes Epsilon, Gamma, Theta, Zeta, Delta and Eta modeled in AF3, Chai-1 and RF2NA.

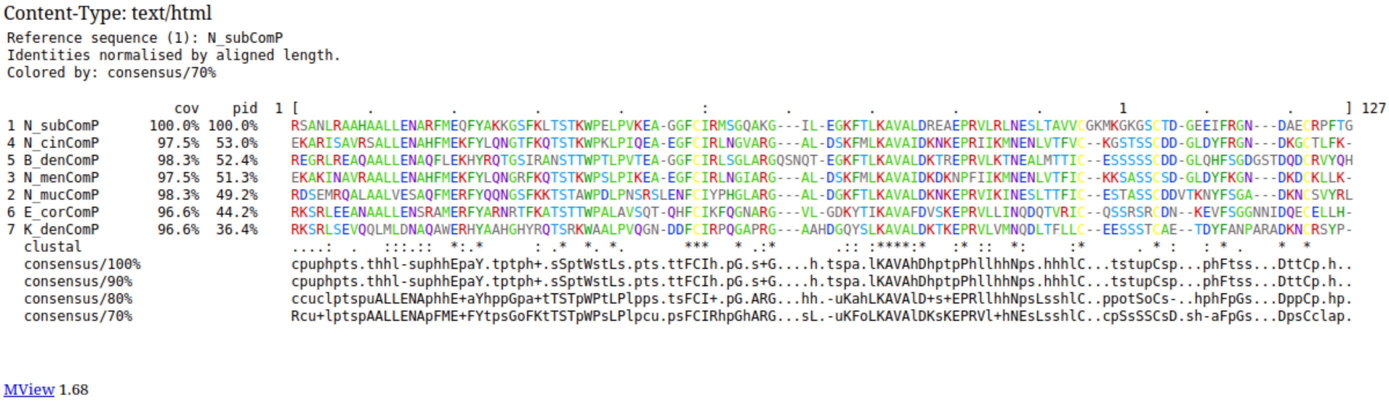

Content-Type: text/html

```
Reference sequence (1): N_subComP
Identities normalised by aligned length.
Colored by: consensus/70%

                    cov     pid  1 [        .         .         .         .         :         .         .         .         .         1         .         .       ] 127
1 N_subComP      100.0% 100.0%    RSANLRAAHAALLENARFMEQFYAKKGSFKLTSTKWPELPVKEA-GGFCIRMSGQAKG---IL-EGKFTLKAVALDREAEPRVLRLNESLTAVVCGKMKGKGSCTD-GEEIFRGN---DAECRPFTG
4 N_cinComP       97.5%  53.0%    EKARISAVRSALLENAHFMEKFYLQNGTFKQTSTKWPKLPIQEA-EGFCIRLNGVARG---AL-DSKFMLKAVAIDKNKEPRIIKMNENLVTFVC--KGSTSSCDD-GLDYFRGN---DKGCTLFK-
5 B_denComP       98.3%  52.4%    REGRLREAQAALLENAQFLEKHYRQTGSIRANSTTWPTLPVTEA-GGFCIRLSGLARGQSNQT-EGKFTLKAVALDKTREPRVLKTNEALMTTIC--ESSSSSCDD-GLQHFSGDGSTDQDCRVYQH
3 N_menComP       97.5%  51.3%    EKAKINAVRAALLENAHFMEKFYLQNGRFKQTSTKWPSLPIKEA-EGFCIRLNGIARG---AL-DSKFMLKAVAIDKDKNPFIIKMNENLVTFIC--KKSASSCSD-GLDYFKGN---DKDCKLLK-
2 N_mucComP       98.3%  49.2%    RDSEMRQALAALVESAQFMERFYQQNGSFKKTSTAWPDLPNSRSLENFCIYPHGLARG---AL-DGKFTLKAVALDKNKEPRVIKINESLTTFIC--ESTASSCDDVTKNYFSGA---DKNCSVYRL
6 E_corComP       96.6%  44.2%    RKSRLEEANAALLENSRAMERFYARNRTFKATSTTWPALAVSQT-QHFCIKFQGNARG---VL-GDKYTIKAVAFDVSKEPRVLLINQDQTVRIC--QSSRSRCDN--KEVFSGGNNIDQECELLH-
7 K_denComP       96.6%  36.4%    RKSRLSEVQQLMLDNAQAWERHYAAHGHYRQTSRKWAALPVQGN-DDFCIRPQGAPRG---AAHDGQYSLKAVALDKTKEPRVLVMNQDLTFLLC--EESSSTCAE--TDYFANPARADKNCRSYP-
  clustal                         ....:  .   :::.::  *:.*      : .*  *. *.       ***   * .:*       .:: :****:*   :* ::  *:      :*       . * :    : * .     *  *
  consensus/100%                  cpuphpts.thhl-suphhEpaY.tptph+.sSptWstLs.pts.ttFCIh.pG.s+G....h.tspa.lKAVAhDhptpPhllhhNps.hhhlC...tstupCsp...phFtss...DttCp.h..
  consensus/90%                   cpuphpts.thhl-suphhEpaY.tptph+.sSptWstLs.pts.ttFCIh.pG.s+G....h.tspa.lKAVAhDhptpPhllhhNps.hhhlC...tstupCsp...phFtss...DttCp.h..
  consensus/80%                   ccuclptspuALLENAphhE+aYhppGpa+tTSTpWPtLPlpps.tsFCI+.pG.ARG...hh.-uKahLKAVAlD+s+EPRllhhNpsLsshlC..ppotSoCs-..hphFpGs...DppCp.hp.
  consensus/70%                   Rcu+lptspAALLENApFME+FYtpsGoFKtTSTpWPsLPlpcu.psFCIRhpGhARG...sL.-uKFoLKAVAlDKsKEPRVl+hNEsLsshlC..cpSsSSCsD.sh-aFpGs...DpsCclap.
```

MView 1.68

**Figure S2.** Alignment of the ComPs, using default settings in MAFFT v. 7.505, visualized in MVIEW v. 1.68. First 28 amino acids of the N. meningitidis mature ComP are trimmed and the other ComPs accordingly.

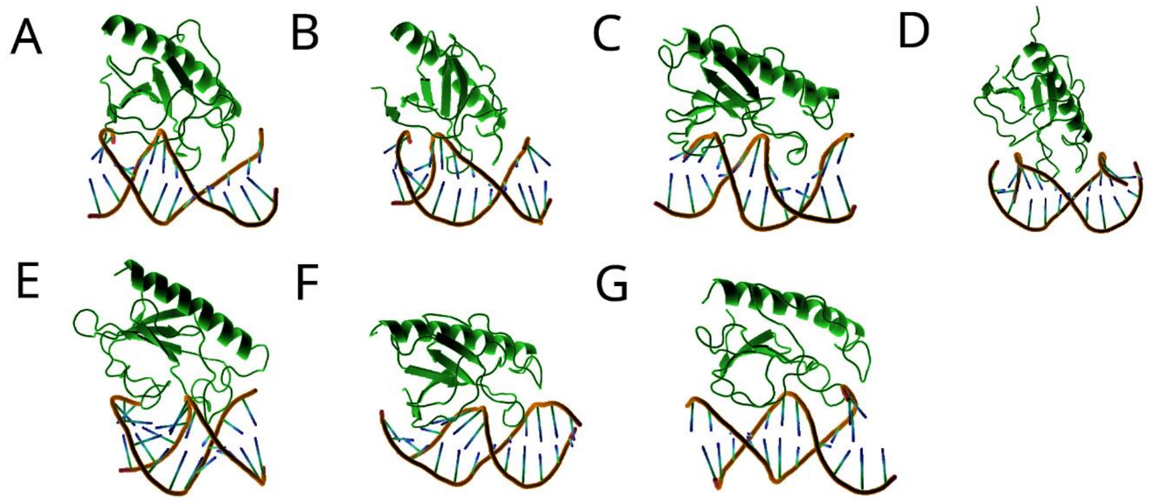


**Figure S3.** Selected high-ranking HADDOCK models, representative of the most abundant binding mode of each ComP$_{nat}$. A: N. muc$_{nat}$ (Epsilon). B: N. men$_{nat}$ (Epsilon). C: K. den$_{nat}$ (Epsilon). D: E. cor$_{nat}$ (Epsilon). E: B. den$_{nat}$ (Epsilon). F: N. cin$_{nat}$ (Gamma). G: N. sub$_{nat}$ (Gamma). AF3, Chai-1 and RF2NA DNA binding modes were used to guide the assignment of grooves in the HADDOCK distorted DNAs.

Figure S4. Representative PAE plots of the top ranking native AF3 model for each $ComP_{nat}$ and assigned mode.

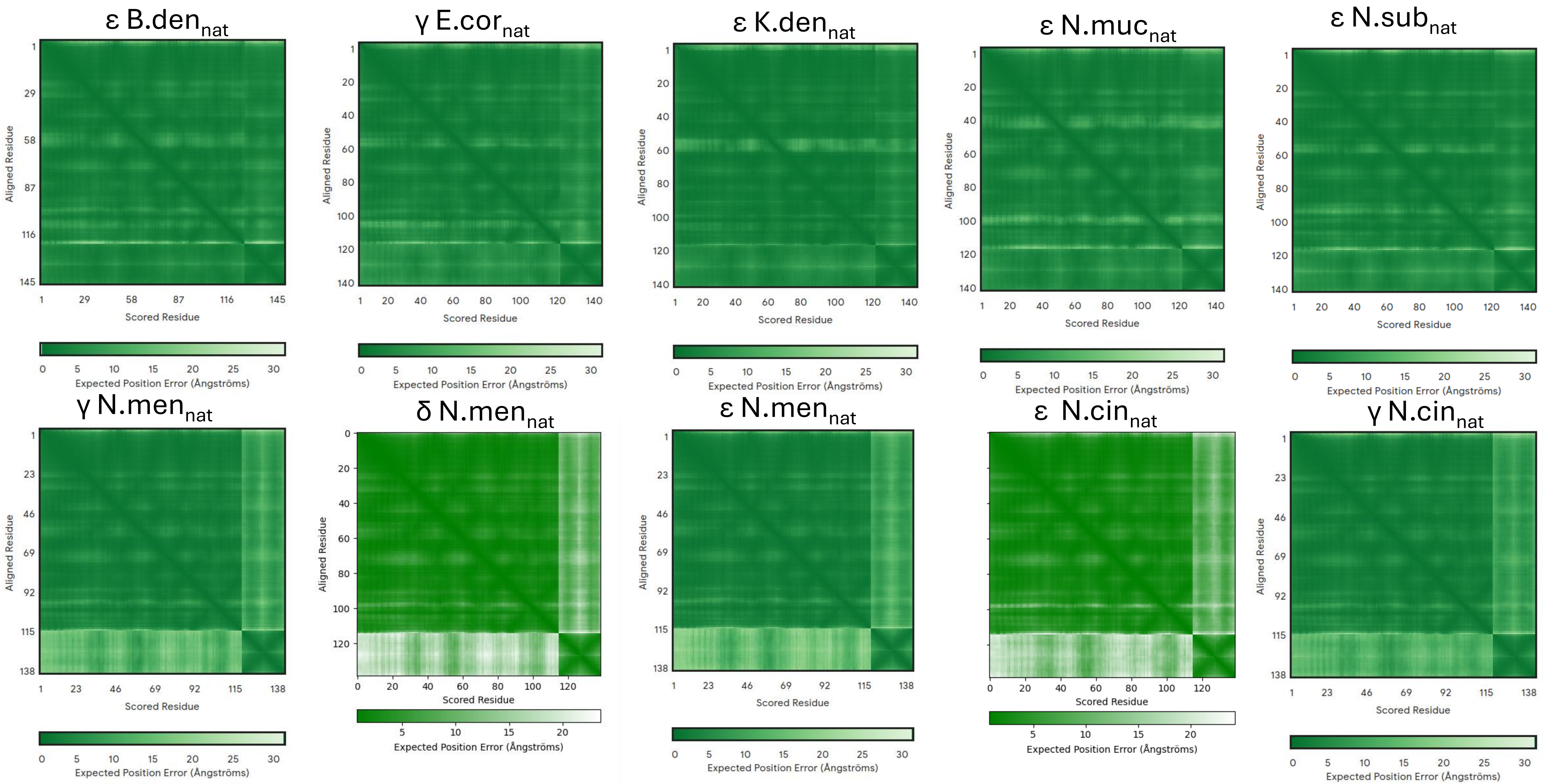

Figure S5. Representative PAE plots of the top ranking native Chai-1 model for each $ComP_{nat}$ and assigned mode.

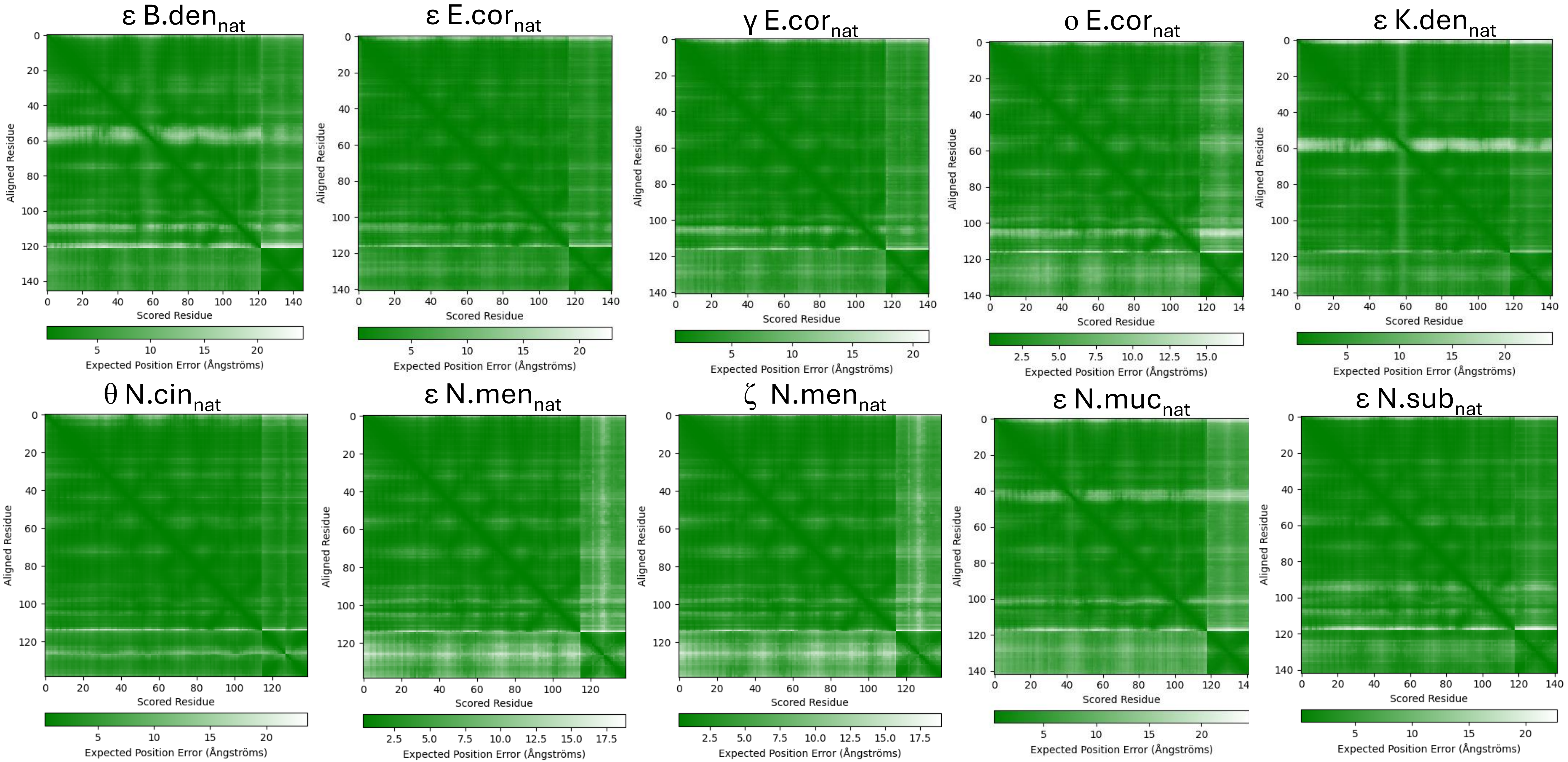

Figure S6. Representative PAE plots of the top ranking native RF2NA model for each ComP$_{nat}$ and assigned mode.

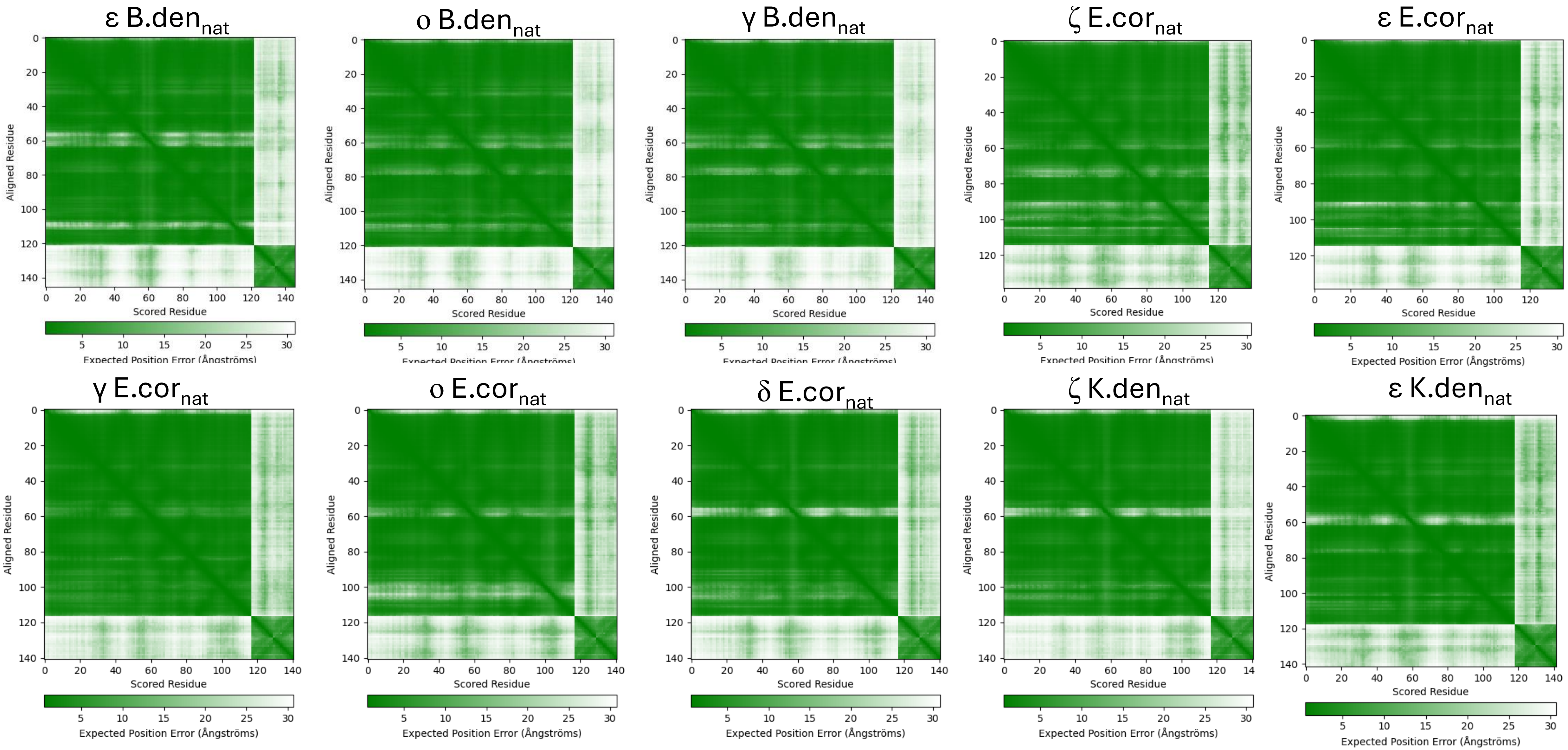

Figure S6. continued

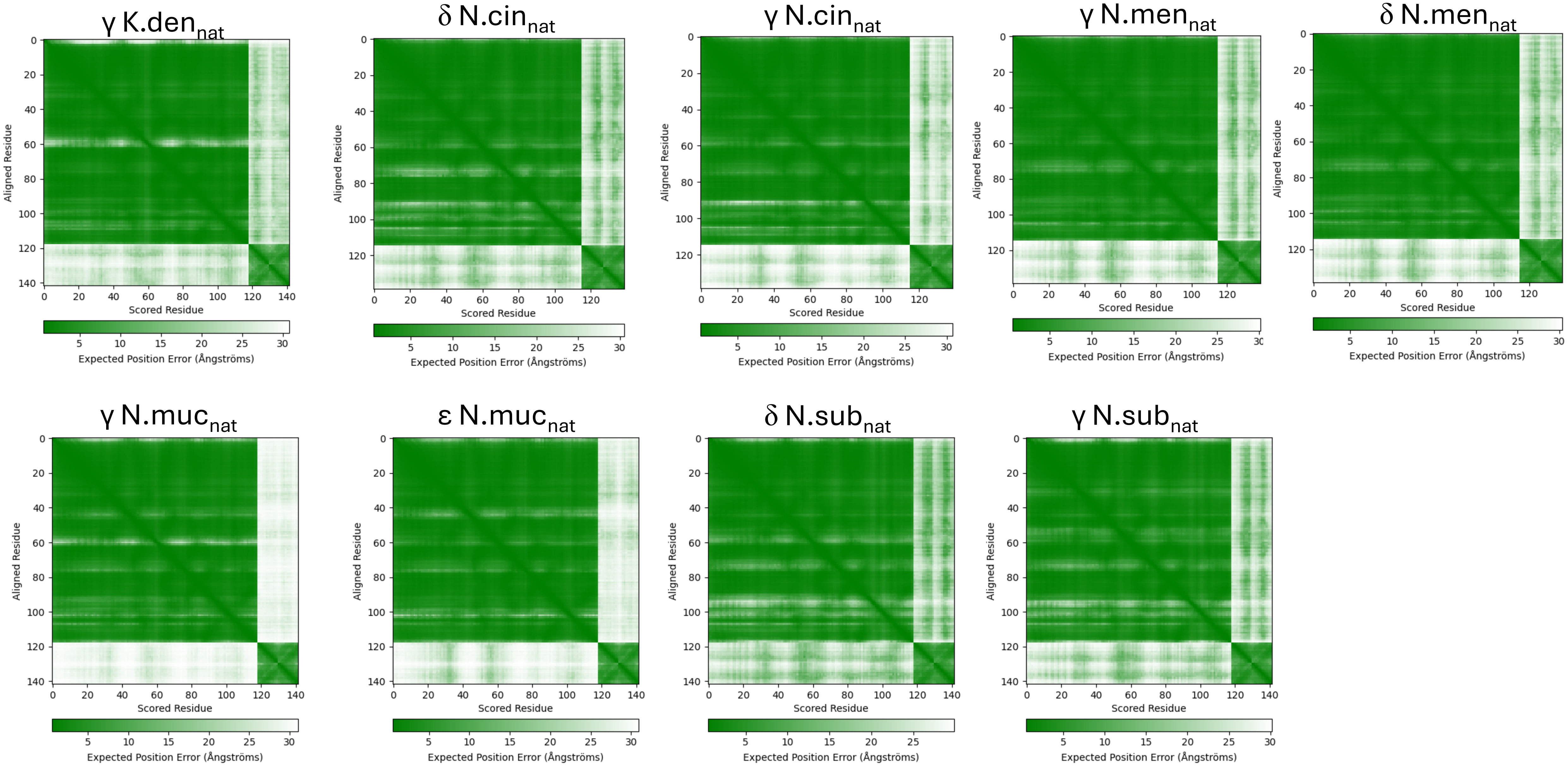

Figure S7. Representative PAE plots of the top ranking scrambled AF3 model for each $ComP_{scr}$ and assigned mode.

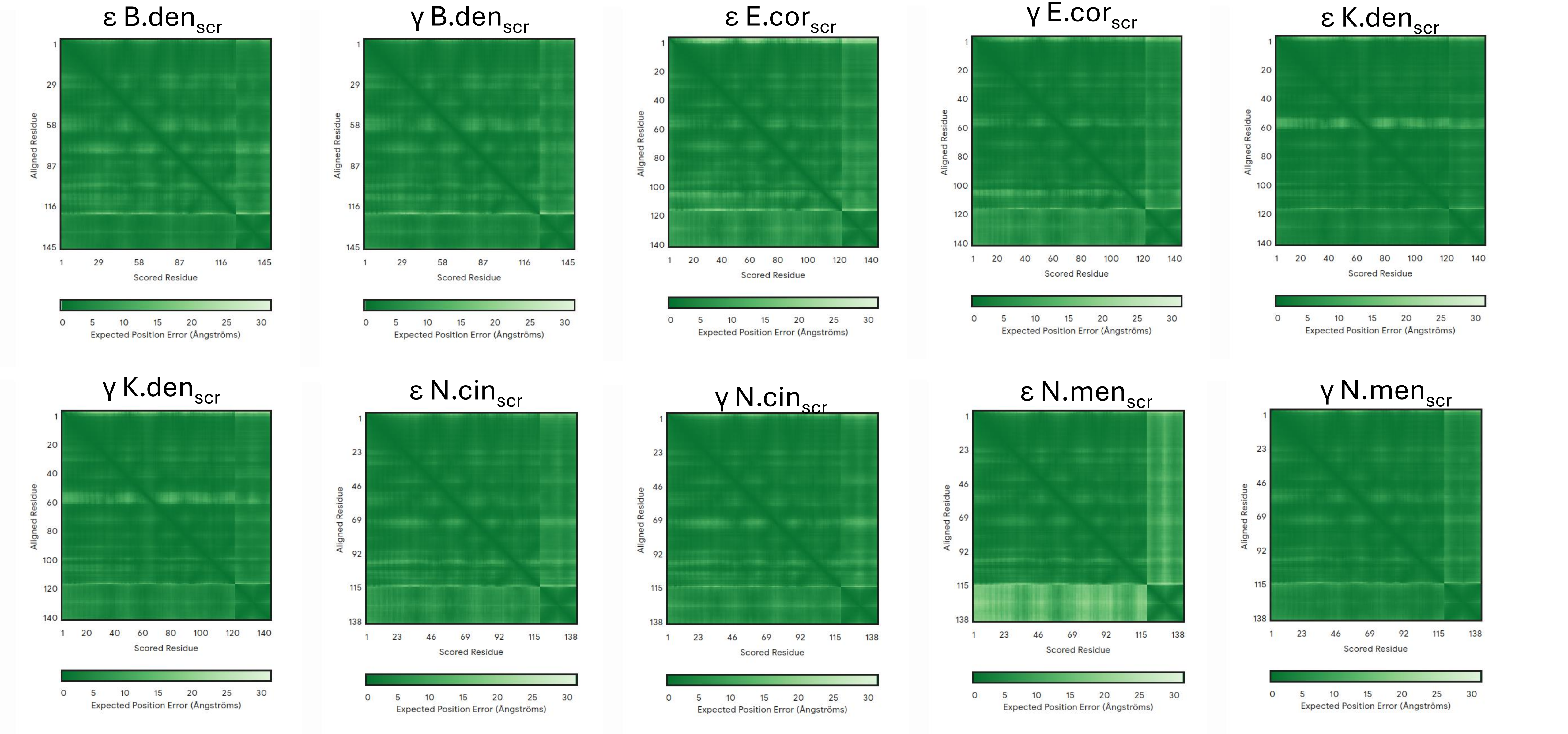

### Figure S7. continued

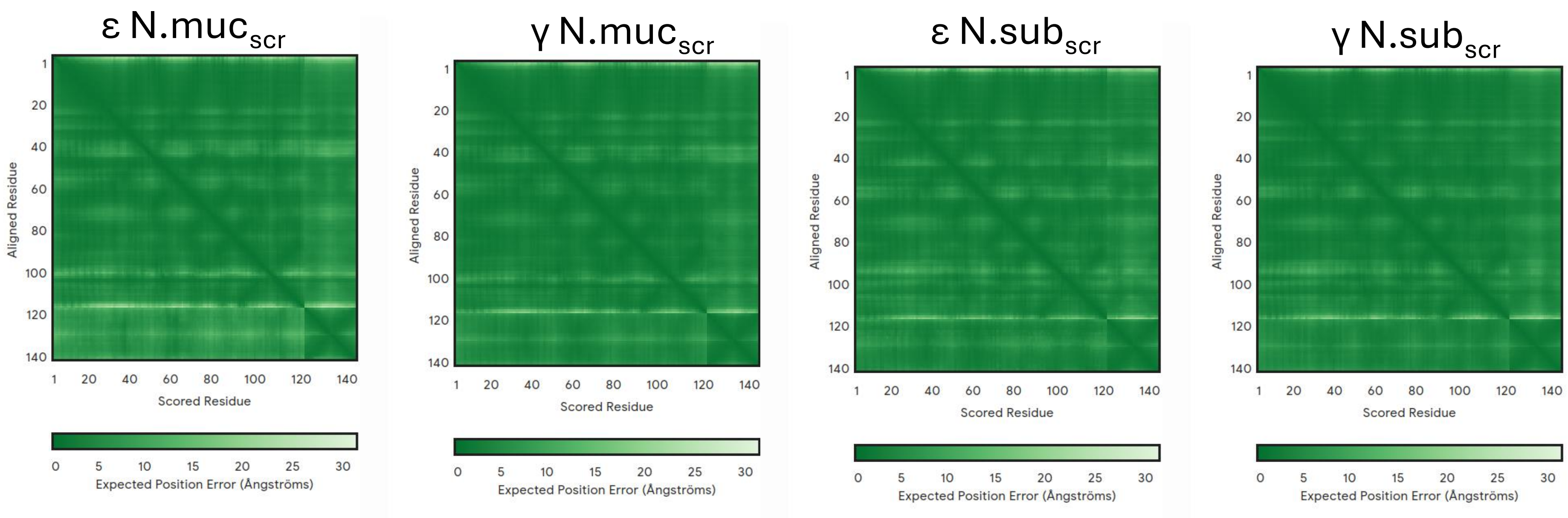

**Code Output S1** Wilcoxon rank-sum tests on PAE and pLDDT, comparing AF3, Chai-1 and RF2NA. Median PAE and pLDDT values are also shown.

```
Species: B_denitrificans
Pairwise Wilcoxon test for PAE:

        Pairwise comparisons using Wilcoxon rank sum test with continuity
correction

data:  PAE and Software

       AF3     Chai-1
Chai-1 0.019   -
RF2NA  8.3e-14 1.2e-12

P value adjustment method: bonferroni
Pairwise Wilcoxon test for pLDDT:

        Pairwise comparisons using Wilcoxon rank sum test with continuity
correction

data:  plddt and Software

       AF3     Chai-1
Chai-1 0.92    -
RF2NA  4.2e-13 1.2e-12

P value adjustment method: bonferroni

Species: E_corrodens
Pairwise Wilcoxon test for PAE:

        Pairwise comparisons using Wilcoxon rank sum test with continuity
correction

data:  PAE and Software

       AF3    Chai-1
Chai-1 <2e-16 -
RF2NA  <2e-16 <2e-16

P value adjustment method: bonferroni
Pairwise Wilcoxon test for pLDDT:

        Pairwise comparisons using Wilcoxon rank sum test with continuity
correction

data:  plddt and Software

       AF3    Chai-1
```

```
Chai-1 <2e-16 -
RF2NA  <2e-16 <2e-16

P value adjustment method: bonferroni

Species: K_denitrificans
Pairwise Wilcoxon test for PAE:

        Pairwise comparisons using Wilcoxon rank sum test with continuity
correction

data:  PAE and Software

       AF3    Chai-1
Chai-1 1      -
RF2NA  <2e-16 <2e-16

P value adjustment method: bonferroni
Pairwise Wilcoxon test for pLDDT:

        Pairwise comparisons using Wilcoxon rank sum test with continuity
correction

data:  plddt and Software

       AF3    Chai-1
Chai-1 <2e-16 -
RF2NA  <2e-16 <2e-16

P value adjustment method: bonferroni

Species: N_cinerea
Pairwise Wilcoxon test for PAE:

        Pairwise comparisons using Wilcoxon rank sum test with continuity
correction

data:  PAE and Software

       AF3    Chai-1
Chai-1 <2e-16 -
RF2NA  <2e-16 <2e-16

P value adjustment method: bonferroni
Pairwise Wilcoxon test for pLDDT:

        Pairwise comparisons using Wilcoxon rank sum test with continuity
correction

data:  plddt and Software
```

```
       AF3     Chai-1
Chai-1 < 2e-16 -
RF2NA  1.8e-11 < 2e-16

P value adjustment method: bonferroni

Species: N_meningitidis
Pairwise Wilcoxon test for PAE:

        Pairwise comparisons using Wilcoxon rank sum test with continuity
correction

data:  PAE and Software

       AF3     Chai-1
Chai-1 1.5e-08 -
RF2NA  < 2e-16 1.9e-08

P value adjustment method: bonferroni
Pairwise Wilcoxon test for pLDDT:

        Pairwise comparisons using Wilcoxon rank sum test with continuity
correction

data:  plddt and Software

       AF3     Chai-1
Chai-1 1.5e-08 -
RF2NA  < 2e-16 1.8e-08

P value adjustment method: bonferroni

Species: N_mucosa
Pairwise Wilcoxon test for PAE:

        Pairwise comparisons using Wilcoxon rank sum test with continuity
correction

data:  PAE and Software

       AF3     Chai-1
Chai-1 < 2e-16 -
RF2NA  1.7e-07 4.1e-08

P value adjustment method: bonferroni
Pairwise Wilcoxon test for pLDDT:

        Pairwise comparisons using Wilcoxon rank sum test with continuity
correction

data:  plddt and Software
```

```
       AF3    Chai-1
Chai-1 <2e-16 -
RF2NA  0.0053 1.0000

P value adjustment method: bonferroni

Species: N_subflava
Pairwise Wilcoxon test for PAE:

        Pairwise comparisons using Wilcoxon rank sum test with continuity
correction

data:  PAE and Software

       AF3    Chai-1
Chai-1 <2e-16 -
RF2NA  <2e-16 <2e-16

P value adjustment method: bonferroni
Pairwise Wilcoxon test for pLDDT:

        Pairwise comparisons using Wilcoxon rank sum test with continuity
correction

data:  plddt and Software

       AF3     Chai-1
Chai-1 0.00027 -
RF2NA  < 2e-16 < 2e-16

P value adjustment method: bonferroni
>
> # Calculate median PAE and plddt values by Software for each Species
> median_values <- merged_df %>%
+   group_by(Species, Software) %>%
+   summarise(
+     median_PAE = median(PAE, na.rm = TRUE),
+     median_plddt = median(plddt, na.rm = TRUE),
+     .groups = "drop"
+   )
>
> # Print median values with species labels
> for (species in unique(median_values$Species)) {
+   cat("\nSpecies:", species, "\n")
+   print(median_values %>% filter(Species == species))
+ }

---

### PAE and pLDDT ###
```

```
> # Perform pairwise Wilcoxon test for PAE and plddt with Bonferroni correction
for each Species
> results <- merged_df %>%
+   group_by(Species) %>%
+   summarise(
+     pairwise_pa = list(pairwise.wilcox.test(PAE, Software, p.adjust.method =
"bonferroni")),
+     pairwise_plddt = list(pairwise.wilcox.test(plddt, Software,
p.adjust.method = "bonferroni"))
+   )
>
> # Print results with species labels
> for (i in 1:nrow(results)) {
+   cat("\nSpecies:", results$Species[i], "\n")
+   cat("Pairwise Wilcoxon test for PAE:\n")
+   print(results$pairwise_pa[[i]])
+   cat("Pairwise Wilcoxon test for pLDDT:\n")
+   print(results$pairwise_plddt[[i]])
+ }

Species: B_denitrificans
# A tibble: 3 × 4
  Species         Software median_PAE median_plddt
  <chr>           <chr>         <dbl>        <dbl>
1 B_denitrificans AF3            4.07         86.9
2 B_denitrificans Chai-1         4.14         87.0
3 B_denitrificans RF2NA          9.90         84.0

Species: E_corrodens
# A tibble: 3 × 4
  Species     Software median_PAE median_plddt
  <chr>       <chr>         <dbl>        <dbl>
1 E_corrodens AF3            4.01         85.0
2 E_corrodens Chai-1         2.89         88.1
3 E_corrodens RF2NA          9.66         82.8

Species: K_denitrificans
# A tibble: 3 × 4
  Species         Software median_PAE median_plddt
  <chr>           <chr>         <dbl>        <dbl>
1 K_denitrificans AF3            3.51         90.8
2 K_denitrificans Chai-1         3.52         85.6
3 K_denitrificans RF2NA          9.48         83.4

Species: N_cinerea
# A tibble: 3 × 4
  Species   Software median_PAE median_plddt
  <chr>     <chr>         <dbl>        <dbl>
1 N_cinerea AF3            5.34         82.4
2 N_cinerea Chai-1         3.05         85.4
```

```
3 N_cinerea RF2NA          9.86         83.6

Species: N_meningitidis
# A tibble: 3 × 4
  Species        Software median_PAE median_plddt
  <chr>          <chr>         <dbl>        <dbl>
1 N_meningitidis AF3            5.73         79.9
2 N_meningitidis Chai-1         3.30         86.3
3 N_meningitidis RF2NA          9.52         84.2

Species: N_mucosa
# A tibble: 3 × 4
  Species  Software median_PAE median_plddt
  <chr>    <chr>         <dbl>        <dbl>
1 N_mucosa AF3            4.09         82.4
2 N_mucosa Chai-1         4.72         83.4
3 N_mucosa RF2NA          9.89         83.7

Species: N_subflava
# A tibble: 3 × 4
  Species    Software median_PAE median_plddt
  <chr>      <chr>         <dbl>        <dbl>
1 N_subflava AF3            4.04         85.9
2 N_subflava Chai-1         3.17         87.6
3 N_subflava RF2NA          9.50         82.0
```

**Code Output S2** Wilcoxon rank-sum tests on ipTM and CPPM for each $ComP_{nat}$.

```
### Native vs. Scrambled DUS ###

> ### Fisher's test for ipTM and CPPM differences across Pairing ###
>
> ### ipTM ###
>
> ### AA-king3DUS
>
> # Subset data for 'Native' Pairing
> group_native <- merged_df_filtered$ipTM[merged_df_filtered$DUS ==
"aa_king3dus" &
+                                            merged_df_filtered$Pairing ==
"Native"]
>
> # Subset data for 'Scrambled' Pairing
> group_scrambled <- merged_df_filtered$ipTM[merged_df_filtered$DUS ==
"aa_king3dus" &
+                                               merged_df_filtered$Pairing ==
"Scrambled"]
>
```

```
> # Test for normality in the 'Native' group
> shapiro_native <- shapiro.test(group_native)
>
> # Test for normality in the 'Scrambled' group
> shapiro_scrambled <- shapiro.test(group_scrambled)
>
> # Print the results
> # print(shapiro_native)
> # print(shapiro_scrambled)
>
> aa_king3dus_df <- subset(merged_df_filtered, DUS == 'aa_king3dus')
>
> # Perform Wilcoxon rank-sum test (non-parametric)
> wilcox_test_result <- wilcox.test(ipTM ~ Pairing, data = aa_king3dus_df,
+                                   exact = FALSE)
>
> # Print the results
> print(wilcox_test_result)

        Wilcoxon rank sum test with continuity correction

data:  ipTM by Pairing
W = 5395.5, p-value = 0.7319
alternative hypothesis: true location shift is not equal to 0

>
> ### AG-DUS
>
> # Subset data for 'Native' Pairing
> group_native <- merged_df_filtered$ipTM[merged_df_filtered$DUS == "ag_dus" &
+                                           merged_df_filtered$Pairing ==
"Native"]
>
> # Subset data for 'Scrambled' Pairing
> group_scrambled <- merged_df_filtered$ipTM[merged_df_filtered$DUS == "ag_dus"
&
+                                              merged_df_filtered$Pairing ==
"Scrambled"]
>
> # Test for normality in the 'Native' group
> shapiro_native <- shapiro.test(group_native)
>
> # Test for normality in the 'Scrambled' group
> shapiro_scrambled <- shapiro.test(group_scrambled)
>
> # Print the results
> # print(shapiro_native)
> # print(shapiro_scrambled)
>
> ag_dus_df <- subset(merged_df_filtered, DUS == 'ag_dus')
>
```

```
> # Perform Wilcoxon rank-sum test (non-parametric)
> wilcox_test_result <- wilcox.test(ipTM ~ Pairing, data = ag_dus_df,
+                                   exact = FALSE, alternative = "greater")
>
> # Print the results
> print(wilcox_test_result)

	Wilcoxon rank sum test with continuity correction

data:  ipTM by Pairing
W = 115384, p-value < 2.2e-16
alternative hypothesis: true location shift is greater than 0

>
> ### AG-eikDUS
>
> # Subset data for 'Native' Pairing
> group_native <- merged_df_filtered$ipTM[merged_df_filtered$DUS == "ag_eikdus"
&
+                                           merged_df_filtered$Pairing ==
"Native"]
>
> # Subset data for 'Scrambled' Pairing
> group_scrambled <- merged_df_filtered$ipTM[merged_df_filtered$DUS ==
"ag_eikdus" &
+                                              merged_df_filtered$Pairing ==
"Scrambled"]
>
> # Test for normality in the 'Native' group
> shapiro_native <- shapiro.test(group_native)
>
> # Test for normality in the 'Scrambled' group
> shapiro_scrambled <- shapiro.test(group_scrambled)
>
> # Print the results
> # print(shapiro_native)
> # print(shapiro_scrambled)
>
> ag_eikdus_df <- subset(merged_df_filtered, DUS == 'ag_eikdus')
>
> # Perform Wilcoxon rank-sum test (non-parametric)
> wilcox_test_result <- wilcox.test(ipTM ~ Pairing, data = ag_eikdus_df,
+                                   exact = FALSE, alternative = "greater")
>
> # Print the results
> print(wilcox_test_result)

	Wilcoxon rank sum test with continuity correction

data:  ipTM by Pairing
W = 8867.5, p-value < 2.2e-16
```

```
alternative hypothesis: true location shift is greater than 0

>
> ### AG-mucDUS
>
> # Subset data for 'Native' Pairing
> group_native <- merged_df_filtered$ipTM[merged_df_filtered$DUS == "ag_mucdus"
&
+                                          merged_df_filtered$Pairing ==
"Native"]
>
> # Subset data for 'Scrambled' Pairing
> group_scrambled <- merged_df_filtered$ipTM[merged_df_filtered$DUS ==
"ag_mucdus" &
+                                             merged_df_filtered$Pairing ==
"Scrambled"]
>
> # Test for normality in the 'Native' group
> shapiro_native <- shapiro.test(group_native)
>
> # Test for normality in the 'Scrambled' group
> shapiro_scrambled <- shapiro.test(group_scrambled)
>
> # Print the results
> # print(shapiro_native)
> # print(shapiro_scrambled)
>
> ag_mucdus_df <- subset(merged_df_filtered, DUS == 'ag_mucdus')
>
> # Perform Wilcoxon rank-sum test (non-parametric)
> wilcox_test_result <- wilcox.test(ipTM ~ Pairing, data = ag_mucdus_df,
+                                    exact = FALSE, alternative = "greater")
>
> # Print the results
> print(wilcox_test_result)

        Wilcoxon rank sum test with continuity correction

data:  ipTM by Pairing
W = 9364, p-value < 2.2e-16
alternative hypothesis: true location shift is greater than 0

>
> ### AT-DUS
>
> # Subset data for 'Native' Pairing
> group_native <- merged_df_filtered$ipTM[merged_df_filtered$DUS == "at_dus" &
+                                          merged_df_filtered$Pairing ==
"Native"]
>
> # Subset data for 'Scrambled' Pairing
```

```
> group_scrambled <- merged_df_filtered$ipTM[merged_df_filtered$DUS == "at_dus"
&
+                                             merged_df_filtered$Pairing ==
"Scrambled"]
>
> # Test for normality in the 'Native' group
> shapiro_native <- shapiro.test(group_native)
>
> # Test for normality in the 'Scrambled' group
> shapiro_scrambled <- shapiro.test(group_scrambled)
>
> # Print the results
> # print(shapiro_native)
> # print(shapiro_scrambled)
>
> at_dus_df <- subset(merged_df_filtered, DUS == 'at_dus')
>
> # Perform Wilcoxon rank-sum test (non-parametric)
> wilcox_test_result <- wilcox.test(ipTM ~ Pairing, data = at_dus_df,
+                                   exact = FALSE, alternative = "less")
>
> # Print the results
> print(wilcox_test_result)

	Wilcoxon rank sum test with continuity correction

data:  ipTM by Pairing
W = 7496, p-value < 2.2e-16
alternative hypothesis: true location shift is less than 0

>
> ### CPPM ###
>
> ### AA-king3DUS
>
> # Subset data for 'Native' Pairing
> group_native <- merged_df_filtered$Chain_pair_pae_min[merged_df_filtered$DUS
== "aa_king3dus" &
+
merged_df_filtered$Pairing == "Native"]
>
> # Subset data for 'Scrambled' Pairing
> group_scrambled <-
merged_df_filtered$Chain_pair_pae_min[merged_df_filtered$DUS == "aa_king3dus" &
+
merged_df_filtered$Pairing == "Scrambled"]
>
> # Test for normality in the 'Native' group
> shapiro_native <- shapiro.test(group_native)
>
> # Test for normality in the 'Scrambled' group
```

```
> shapiro_scrambled <- shapiro.test(group_scrambled)
>
> # Print the results
> # print(shapiro_native)
> # print(shapiro_scrambled)
>
> aa_king3dus_df <- subset(merged_df_filtered, DUS == 'aa_king3dus')
>
> # Perform Wilcoxon rank-sum test (non-parametric)
> wilcox_test_result <- wilcox.test(Chain_pair_pae_min ~ Pairing, data =
aa_king3dus_df,
+                                   exact = FALSE)
>
> # Print the results
> print(wilcox_test_result)

        Wilcoxon rank sum test with continuity correction

data:  Chain_pair_pae_min by Pairing
W = 5884.5, p-value = 0.1353
alternative hypothesis: true location shift is not equal to 0

>
> ### AG-DUS
>
> # Subset data for 'Native' Pairing
> group_native <- merged_df_filtered$Chain_pair_pae_min[merged_df_filtered$DUS
== "ag_dus" &
+
merged_df_filtered$Pairing == "Native"]
>
> # Subset data for 'Scrambled' Pairing
> group_scrambled <-
merged_df_filtered$Chain_pair_pae_min[merged_df_filtered$DUS == "ag_dus" &
+
merged_df_filtered$Pairing == "Scrambled"]
>
> # Test for normality in the 'Native' group
> shapiro_native <- shapiro.test(group_native)
>
> # Test for normality in the 'Scrambled' group
> shapiro_scrambled <- shapiro.test(group_scrambled)
>
> # Print the results
> # print(shapiro_native)
> # print(shapiro_scrambled)
>
> ag_dus_df <- subset(merged_df_filtered, DUS == 'ag_dus')
>
> # Perform Wilcoxon rank-sum test (non-parametric)
> wilcox_test_result <- wilcox.test(Chain_pair_pae_min ~ Pairing, data =
```

```
ag_dus_df,
+                                   exact = FALSE, alternative = "less")
>
> # Print the results
> print(wilcox_test_result)

        Wilcoxon rank sum test with continuity correction

data:  Chain_pair_pae_min by Pairing
W = 18604, p-value < 2.2e-16
alternative hypothesis: true location shift is less than 0

>
> ### AG-eikDUS
>
> # Subset data for 'Native' Pairing
> group_native <- merged_df_filtered$Chain_pair_pae_min[merged_df_filtered$DUS
== "ag_eikdus" &
+
merged_df_filtered$Pairing == "Native"]
>
> # Subset data for 'Scrambled' Pairing
> group_scrambled <-
merged_df_filtered$Chain_pair_pae_min[merged_df_filtered$DUS == "ag_eikdus" &
+
merged_df_filtered$Pairing == "Scrambled"]
>
> # Test for normality in the 'Native' group
> shapiro_native <- shapiro.test(group_native)
>
> # Test for normality in the 'Scrambled' group
> shapiro_scrambled <- shapiro.test(group_scrambled)
>
> # Print the results
> # print(shapiro_native)
> # print(shapiro_scrambled)
>
> ag_eikdus_df <- subset(merged_df_filtered, DUS == 'ag_eikdus')
>
> # Perform Wilcoxon rank-sum test (non-parametric)
> wilcox_test_result <- wilcox.test(Chain_pair_pae_min ~ Pairing, data =
ag_eikdus_df,
+                                   exact = FALSE, alternative = "less")
>
> # Print the results
> print(wilcox_test_result)

        Wilcoxon rank sum test with continuity correction

data:  Chain_pair_pae_min by Pairing
W = 1840, p-value = 5.809e-15
```

```
alternative hypothesis: true location shift is less than 0

>
> ### AG-mucDUS
>
> # Subset data for 'Native' Pairing
> group_native <- merged_df_filtered$Chain_pair_pae_min[merged_df_filtered$DUS
== "ag_mucdus" &
+
merged_df_filtered$Pairing == "Native"]
>
> # Subset data for 'Scrambled' Pairing
> group_scrambled <-
merged_df_filtered$Chain_pair_pae_min[merged_df_filtered$DUS == "ag_mucdus" &
+
merged_df_filtered$Pairing == "Scrambled"]
>
> # Test for normality in the 'Native' group
> shapiro_native <- shapiro.test(group_native)
>
> # Test for normality in the 'Scrambled' group
> shapiro_scrambled <- shapiro.test(group_scrambled)
>
> # Print the results
> # print(shapiro_native)
> # print(shapiro_scrambled)
>
> ag_mucdus_df <- subset(merged_df_filtered, DUS == 'ag_mucdus')
>
> # Perform Wilcoxon rank-sum test (non-parametric)
> wilcox_test_result <- wilcox.test(Chain_pair_pae_min ~ Pairing, data =
ag_mucdus_df,
+                                    exact = FALSE, alternative = "less")
>
> # Print the results
> print(wilcox_test_result)

        Wilcoxon rank sum test with continuity correction

data:  Chain_pair_pae_min by Pairing
W = 1200, p-value < 2.2e-16
alternative hypothesis: true location shift is less than 0

>
> ### AT-DUS
>
> # Subset data for 'Native' Pairing
> group_native <- merged_df_filtered$Chain_pair_pae_min[merged_df_filtered$DUS
== "at_dus" &
+
merged_df_filtered$Pairing == "Native"]
```

```
>
> # Subset data for 'Scrambled' Pairing
> group_scrambled <-
merged_df_filtered$Chain_pair_pae_min[merged_df_filtered$DUS == "at_dus" &
+
merged_df_filtered$Pairing == "Scrambled"]
>
> # Test for normality in the 'Native' group
> shapiro_native <- shapiro.test(group_native)
>
> # Test for normality in the 'Scrambled' group
> shapiro_scrambled <- shapiro.test(group_scrambled)
>
> # Print the results
> # print(shapiro_native)
> # print(shapiro_scrambled)
>
> at_dus_df <- subset(merged_df_filtered, DUS == 'at_dus')
>
> # Perform Wilcoxon rank-sum test (non-parametric)
> wilcox_test_result <- wilcox.test(Chain_pair_pae_min ~ Pairing, data =
at_dus_df,
+                                    exact = FALSE, alternative = "greater")
>
> # Print the results
> print(wilcox_test_result)

        Wilcoxon rank sum test with continuity correction

data:  Chain_pair_pae_min by Pairing
W = 32456, p-value < 2.2e-16
alternative hypothesis: true location shift is greater than 0
```

**Code Output S3** Wilcoxon rank-sum test on DockQ scores for the internal platform consistency check.

```
> # Display results
> test_af3_chai

      Wilcoxon rank sum test with continuity correction

data:  af3_df$DockQ and chai_df$DockQ
W = 1.3061e+10, p-value < 2.2e-16
alternative hypothesis: true location shift is not equal to 0

> test_af3_rf2na

      Wilcoxon rank sum test with continuity correction
```

```
data:  af3_df$DockQ and rf2na_df$DockQ
W = 432365430, p-value < 2.2e-16
alternative hypothesis: true location shift is not equal to 0

> test_chai_rf2na

	Wilcoxon rank sum test with continuity correction

data:  chai_df$DockQ and rf2na_df$DockQ
W = 3699847036, p-value < 2.2e-16
alternative hypothesis: true location shift is not equal to 0

>
> median(af3_df$DockQ)
[1] 0.841
> median(chai_df$DockQ)
[1] 0.68
> median(rf2na_df$DockQ)
[1] 0.617
```
